# Modulation of CIP2A dimerization and protein stability via druggable pocket

**DOI:** 10.1101/2025.10.13.682046

**Authors:** Karolina Pavic, Paola Moyano-Gómez, Pekka Roivas, Sami T. Kurkinen, Hanna Parkkola, Linda Kauppinen, Sanna Rauhamäki, Aleksi Lehtinen, Petja Rosenqvist, Pasi Virta, Ulla Pentikäinen, Olli T. Pentikäinen, Jukka Westermarck

## Abstract

Cancerous inhibitor of PP2A (CIP2A) supports malignant growth across several cancer types. CIP2A is also a synthetic lethal therapy target for BRCA-mutant cancers. In addition, CIP2A causes Alzheimeŕs disease (AD) phenotype in mice. However, it is not dispensable for normal development and growth. Therefore, CIP2A is very lucrative therapy target in cancer and AD. So far it has been unclear whether the CIP2A protein harbors any druggable pockets amenable to targeting by small molecules.

Here we discover a druggable pocket adjacent to the homodimerization domain of CIP2A. Mutagenesis demonstrates that pocket impacts CIP2A homodimerization, PP2A interaction, and full-length CIP2A protein stability in cancer cells. Further, we identify Gambogenic acid (GNA) as a tool compound that directly binds to CIP2A pocket region. GNA and its derivative Gambogic acid dissociate CIP2A homodimer *in vitro and* cause CIP2A destabilization *in cellulo*. The pocket contributes to the impact of GNA on homodimerization based on the protein fragment and mutagenesis analysis.

These results identify structural vulnerability on CIP2A and may facilitate development of targeted CIP2A inhibitors with broad disease applicability.

## Introduction

Protein phosphatase 2A (PP2A) complexes are widely involved in regulation of cellular serine/ threonine phosphorylation and thereby contribute to basically all physiological processes (Reynhout & Janssens, 2019). When dysregulated, PP2A-mediated protein dephosphorylation becomes pathogenic and leads to many common diseases such as cancer, Alzheimeŕs disease (AD), immune disorders, and diabetes. Therefore, pharmacological targeting of PP2Ás activity could offer totally new opportunities for disease control in common human diseases with unmet medical needs. However, only recently has our understanding of structural mechanisms of PP2A complex regulation reached the level that has facilitated first therapeutic approaches targeting PP2A. (Vainonen, Momeny et al., 2021).

Some PP2A complexes, particularly those containing B56 (PPP2R5) subunits function as tumor suppressors by their capability to dephosphorylate several oncoproteins such as MYC (Fowle, Zhao et al., 2019). The most prevalent mechanism by which PP2A-B56 is inhibited in human cancers is overexpression of an oncogenic inhibitor protein Cancerous Inhibitor of PP2A (CIP2A)(Chen, Hu et al., 2023, Junttila, Puustinen et al., 2007, Nagelli & Westermarck, 2024). CIP2A is an obligate homodimer and dimerization both promotes interaction with B56 and protects CIP2A from protein degradation (Wang, Okkeri et al., 2017). However, the structural mechanisms how CIP2A homodimerization might be regulated have not been addressed so far.

Based on numerous studies, CIP2A overexpression is one of the most common oncogenic mechanisms across human cancer types (Chen et al., 2023, Tang, Shen et al., 2018). Functionally CIP2A overexpression supports anchorage independence, epithelial-mesenchymal transition, *in vivo* tumor growth, and resistance on cancer cells across dozens of cancer therapies in different cancer types (Chen et al., 2023, Kauko, Imanishi et al., 2020, Kauko, O’Connor et al., 2018, Khanna, Pimanda et al., 2013, Nagelli & Westermarck, 2024). In addition, CIP2A was very recently recognized as a synthetic lethal partner in BRCA mutant and homologous recombination defective (HRD) cancers, via its direct interaction with central DNA damage response protein TopBP1 (Adam, Rossi et al., 2021, Laine, Nagelli et al., 2021, Lin, Hu et al., 2023, Nagelli & Westermarck, 2024). Thereby CIP2A is a very prominent cancer therapy target across different cancer types and genomic backgrounds. Its role as a therapy target is strongly supported by the lack of detrimental developmental and growth effects on the CIP2A deficient mouse model, which, however, is impaired in both HER2-driven and DNA damage-induced mammary tumorigenesis (Laine et al., 2021, Laine, Sihto et al., 2013, Ventelä, Côme et al., 2012).

In addition to cancer, CIP2A-mediated PP2A inhibition also has a critical role in AD pathogenesis. Phosphorylated Tau was one of the first recognized PP2A targets and PP2A dephosphorylates another critical AD protein amyloid precursor protein (APP). A critical role for CIP2A-mediated PP2A inhibition in Tau and APP phosphorylation and AD pathogenesis was recently established (Hu, Wang et al., 2022, Shentu, Huo et al., 2018). Exogenous overexpression of CIP2A in a healthy mouse brain induces full-blown mouse AD-like pathogenesis and symptoms associated with synaptic degeneration and cognitive effects (Shentu et al., 2018). In addition to pathogenic effects on glial and neural cells, CIP2A overexpression induces *in vivo* neuroinflammation associated with AD initiation and progression (Zhou, Liu et al., 2022).

Importantly, direct pharmaceutical activation of CIP2Ás target, the B56-PP2A complex, has strong anti-oncogenic activity and ameliorates AD pathogenesis in cell and animal models (Kauko et al., 2018, Leonard, Huang et al., 2020, Wei, Zhang et al., 2020). Therefore B56-PP2A reactivation via CIP2A inhibition could provide a lucrative approach for disease control both in cancer and AD. CIP2A therapies inducing CIP2A protein degradation would not only inhibit CIP2A functions towards PP2A-B56α, but would also remove any other potential activities of the protein such as those mediated by direct CIP2A-TopBP1 binding in HRD cancers (Adam et al., 2021, Laine et al., 2021, Nagelli & Westermarck, 2024). Several molecules have been reported as of today to induce inhibition of CIP2A expression via either transcriptional inhibition or by other yet undefined mechanisms (Chen et al., 2023), but currently, there is no evidence for a small molecule binding pocket on CIP2A. Identification of a small-molecule druggable pocket on CIP2A would constitute a major advancement towards structure-based CIP2A inhibitor development for cancer and AD.

Here we identify a pocket-like structure adjacent to CIP2A homodimerization surface and demonstrate that it has an important structural role in regulation of CIP2A homodimerization and consequently CIP2A protein stability. This increased structural understanding of CIP2A provides critical structural coordinates for development of pocket-targeting pharmacological approaches. Eventually, we identify Gambogenic acid and its close derivatives as tool compounds interacting with the pocket, and impairing CIP2A dimerization and protein stability.

## Results

We analyzed the crystal structure of CIP2A homodimer (PDB: 5UFL)(Wang et al., 2017) to identify potential small-molecule binding pockets using the Panther method, by which atomistic shape-electrostatic models of potential ligand-binding areas in a protein can be created by utilizing the protein crystal structure (Niinivehmas, Salokas et al., 2015). Notably, the search resulted in identification of a suitable ligand binding cavity in the C-terminal end of CIP2A 1-560, located near the dimerization interface (Fig. 1A,B). This cavity is partly shielded from the surrounding solvent by intramolecular hydrogen-bonding interaction (D477-Q524) over the underlining groove (Fig. 1B). Another relevant notion was that the lower part of the pocket was connected to the groove extending to K490 (Fig. 1A,B), which was shown in a recent study to cross-link with the B56 subunit of PP2A (Pavic, Gupta et al., 2023). Notably, this same region was indicated as a high probability small-molecule binding pocket also by P2Rank, a machine learning-based method for prediction of ligand binding sites from protein structure (Fig. 1C)(Krivak & Hoksza, 2018). Very importantly, both prediction programs indicated that Q558 from another monomer of the CIP2A homodimer is part of the pocket (Fig. 1B,C). This suggests both that the formation of the full pocket requires CIP2A homodimerization, but also that targeting of the pocket might interfere with the dimerization. Hereafter, we will refer to this region as the CIP2A pocket.

**Figure 1:**
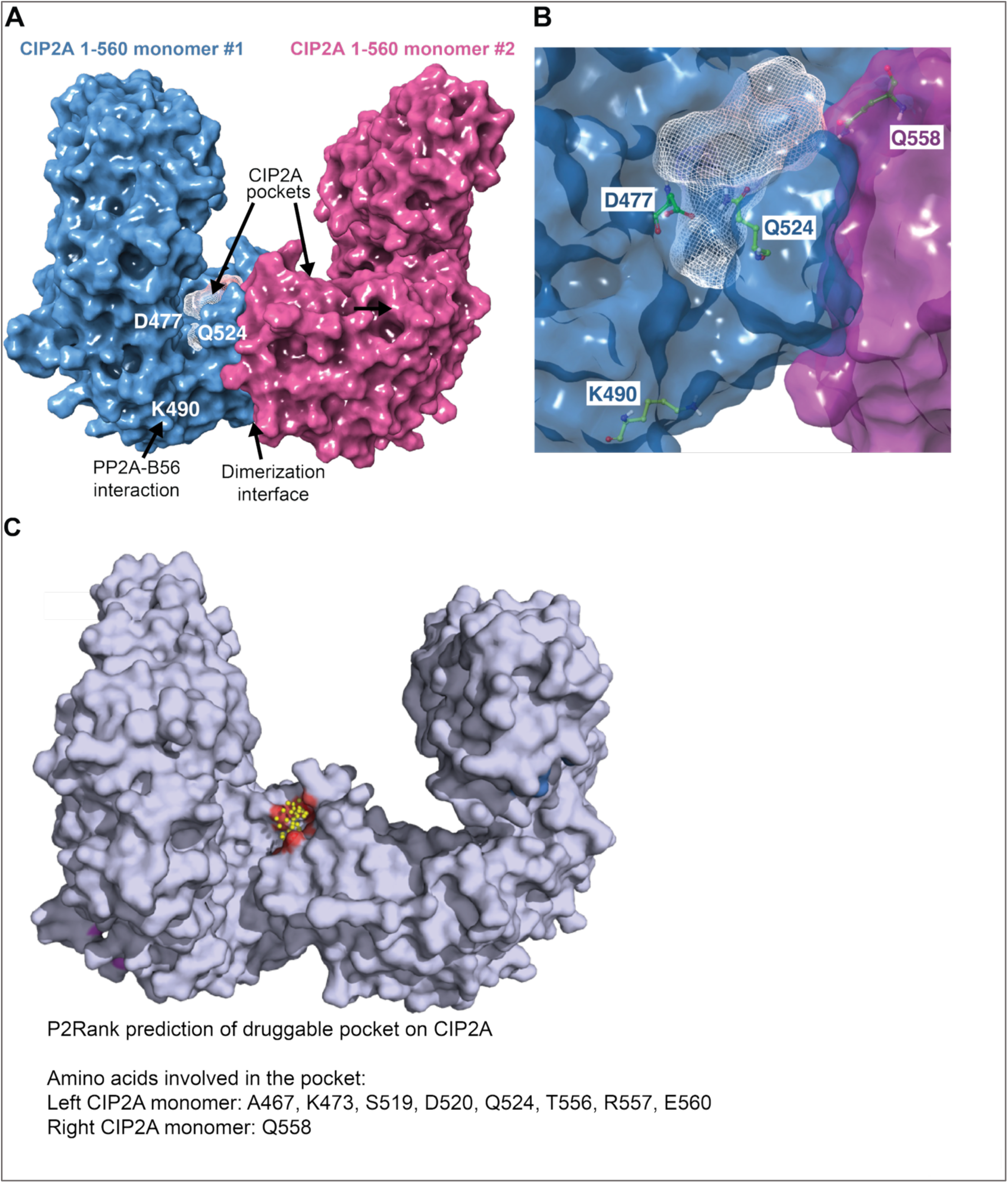
Identification of a pocket-like structure in CIP2A. (**A**) Identification of the pocket-like structure on CIP2A by the Panther method. Mesh indicates the location of the pocket-like structure in the CIP2A(1-560) fragment in proximity to the dimer interface, where the K490 residue was previously identified to be inter-molecularly cross-linked to the regulatory B56α-PP2A subunit. (**B**) Close-up view of the identified CIP2A pocket at the dimer interface, highlighting D477, K490, Q524 and Q558 from the opposite chain. Within this pocket, K490 is prominently positioned, underscoring its structural relevance in CIP2A and its potential role in mediating protein-protein interactions. (**C**) Predicted druggable pockets on CIP2A (PDB ID: 5UFL) identified using the P2Rank machine-learning algorithm. The pocket encompassing residue R577 is highlighted in red. Ligandability score region is indicated by yellow spheres.

### CIP2A pocket is a structural hotspot for the regulation of CIP2A homodimerization and protein stability

Next, we conducted *in silico* exploration of mutations within the CIP2A pocket that could influence CIP2A function. To this end, the alanine to tryptophan mutation was introduced to amino acid 476 (A476W) *in silico,* and molecular dynamics (MD) simulations were performed to understand the potential effects of this mutation. Introducing tryptophan at a key position causes a nearby arginine 557 to shift towards opposing monomer glutamine 558. This reciprocal effect at both monomers weakens the interface, potentially leading to dimer dissociation. (Fig. 2A).

**Figure 2:**
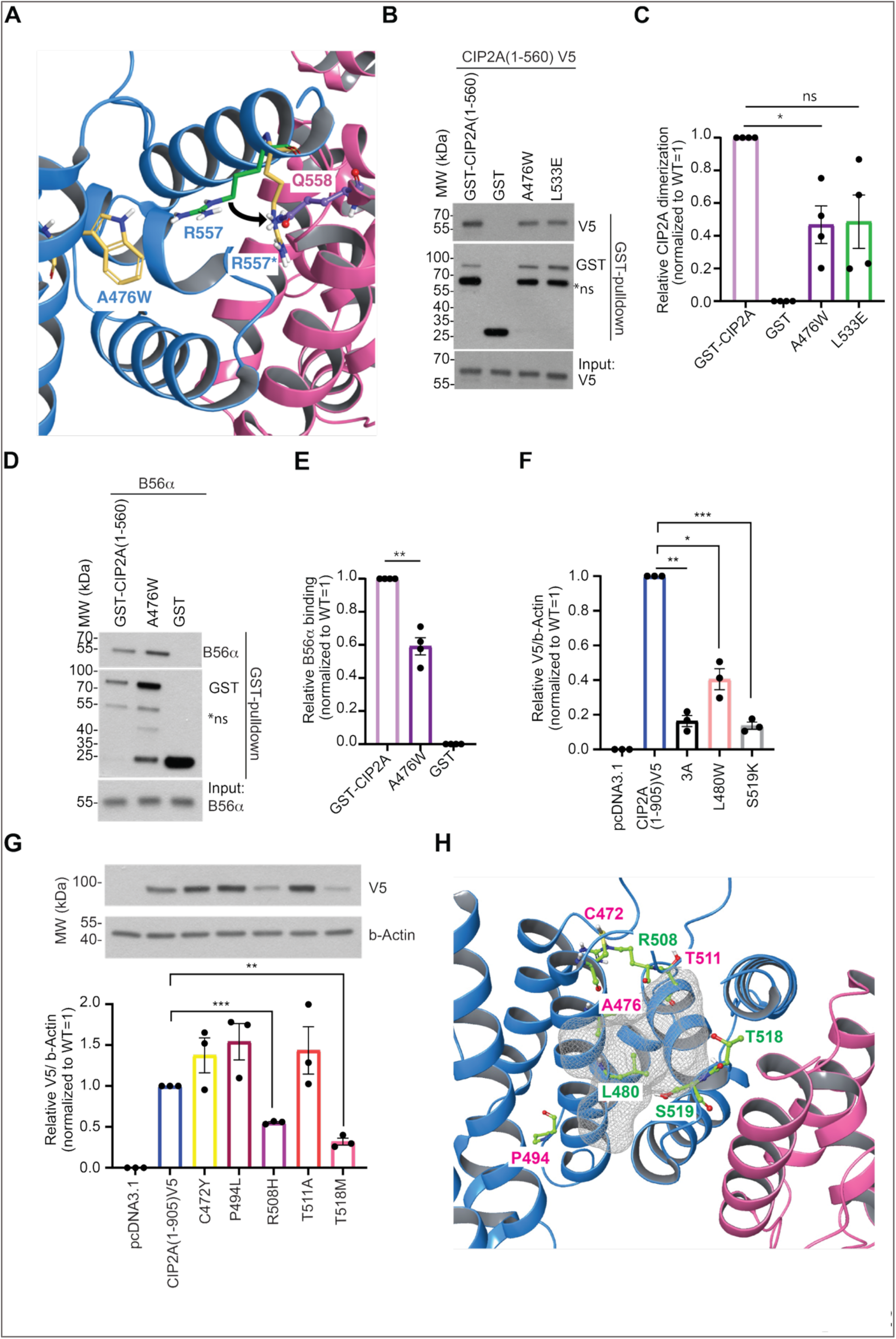
The CIP2A pocket is a structural hotspot for regulation of homodimerization and protein stability. (**A**) Molecular dynamics simulations reveal that A476W substitution (orange sticks) induces a conformational shift of R557 (green sticks, * and yellow stick denotes the shifted R557 position) of the same monomer toward the Q558 (purple sticks) of the other monomer across the dimer interface, thereby destabilizing the CIP2A dimer. (**B,C**) Representative blot of the GST-pull-down showing reduced heterodimerization of the A476W CIP2A mutant in comparison to the dimerization defective L533E mutant (B), with quantified mean ± SEM of N = 3 biological repeats (C). (**D,E**) Representative blot of the GST-pull-down showing reduced B56α of the A476W CIP2A mutant in comparison to the L533E mutant (D), with quantified mean ± SEM of N = 3 biological repeats (E). (**F**) The impact of pocket amino acids on CIP2A full-length CIP2A protein stability in 22RV1 prostate cancer cell. Shown are relative protein expression levels of two pocket mutations L480W and S519K as compared to WT CIP2A(1-905) protein and 3A mutant with triple mutations directly at the dimer interface, with quantified mean ± SEM of N = 3 biological repeats. (**G**) Representative blot showing impact of CIP2A pocket mutations extracted from COSMIC database on CIP2A full-length protein stability as compared to WT CIP2A(1-905) protein with quantified mean ± SEM of N = 3 biological repeats. (**H**) Schematic figure summarizing the stabilizing (green) or destabilizing (red) effect of a subset of tested CIP2A pocket amino acids mapped onto the CIP2A pocket structure (PDB: 5UFL).

These MD results were validated by an *in vitro* dimerization assay using GST- and V5-tagged recombinant CIP2A(1–560) proteins. Consistent with the MD predictions, the A476W pocket mutation reduced CIP2A homodimerization to a level comparable to the previously characterized L533E mutation of the actual dimerization interface (Wang et al., 2017)(Fig. 2B,C). We also tested the impact of the A476W pocket mutation on direct CIP2A-B56a interaction due to the vicinity of the pocket from K490 (Fig. 1A,B), and previous demonstration of the positive impact of homodimerization on the CIP2A-B56α interaction stability (Pavic et al., 2023, Wang et al., 2017). In line with our prediction, the pocket mutant A476W exhibited significantly reduced binding to B56α as compared to GST-CIP2A(1-560) (Fig. 2D,E).

To expand these novel findings indicating important functional role for the CIP2A pocket, we employed our standard *in cellulo* assay in which the impact of CIP2A pocket point is studied by introducing the mutations on V5-tagged full-length CIP2A(1-905) mammalian expression construct, and examination of CIP2Ás protein expression by Western blotting. The chosen strategy was supported by previous data that CIP2A protein stability in cells directly reflects its homodimerization efficiency (Wang et al., 2017). Furthermore, this was the only available approach to study the contribution of the pocket amino acids in the context of full-length CIP2A protein because production of recombinant full-length CIP2A protein, in quantities and qualities required for functional experiments, remains elusive. Notably, out of the tested point mutations, L480W and S519K mutations very efficiently destabilized CIP2A protein expression in cancer cells, as compared to WT protein (Fig. 2F and Fig. S1A). Linking the protein stability effects to inhibition of CIP2A homodimerization, both these pocket mutations induced comparable destabilization effect as triple alanine substitutions L529A/L532A/L533A (3A) of amino acids directly at the homodimerization surface (Fig. 2F, S1B)(Wang et al., 2017).

Last, acknowledging the functional importance of CIP2A protein expression levels for cancer phenotypes, we explored the potential impact of cancer-derived CIP2A pocket mutations on protein expression. For this purpose, we extracted CIP2A gene cancer mutations registered in the COSMIC database. Based on the positioning of the mutated residues and their anticipated effect on pocket structure, we selected five cancer-derived CIP2A pocket mutations for functional testing. Notably, while three of the cancer-derived pocket mutations (C472Y, P494L, T511A) exhibited increased CIP2A protein expression, R508H and T518M had detrimental destabilizing effects on CIP2A expression (Fig. 2G). All CIP2A pocket amino acids identified to have impact either on CIP2A dimerization, B56 interaction and/or full-length protein expression are color-coded (red; increased stability, green; decreased stability) to pocket structure in the Fig. 2H.

As a summary, these results identify the CIP2A pocket as a critical structural feature impacting CIP2A homodimerization and protein stability.

### Small molecule docking of Gambogenic acid to the CIP2A pocket

Intrigued by the importance of the CIP2A pocket for CIP2A function and stability, we questioned whether there would be small molecules that could induce similar structural disturbance by binding to the CIP2A pocket. Several small molecules have been shown to inhibit CIP2A protein expression *in cellulo*, although the exact mechanism of their mode of action, or whether the inhibitory effect was due to direct targeting of CIP2A, often remained poorly understood (Chen et al., 2023). Further, so far, no evidence has been published for pocket-like structures mediating the inhibitory effects of small molecules on CIP2A. Therefore, we mined the literature and extracted compounds proposed to be CIP2A inhibitors. We superposed the proposed CIP2A inhibitors to the pocket by ShaEP method (Vainio, Puranen et al., 2009), and shortlisted gambogenic acid (GNA; CAS 173932-75-7) as a possible binder to the CIP2A pocket. GNA is a natural compound previously shown to induce proteosomal degradation of CIP2A across several cell lines with low micromolar concentration, although no evidence for direct binding of GNA to CIP2A was presented (Yu, Zhao et al., 2016). Docking of GNA to the crystal structure of CIP2A dimer and monomer enabled us to identify possible binding conformation that could interfere with the Q558 position from the other monomer, and thus potentially affect the dimerization of CIP2A (Fig. 3A,B). Importantly, using microscale thermophoresis (MST), GNA was found to directly bind to CIP2A(1-560), with the dissociation constant K_D_ of 780 nM (Fig. 3C). Furthermore, we performed literature search for similar compound scaffolds and identified gambogic acid (GIA; CAS 2752-65-0) and DAP-19 (Yim, Prince et al., 2016)(Fig. S2A). Whereas GIA has a longer hydrophobic tail than gambogic acid and DAP-19, due to the opening of its pyrano ring, in DAP-19 the carboxylic acid of GIA has been converted to a morpholine amide (Fig. S2A)(Yim et al., 2016). Both GIA and DAP-19 bound to CIP2A(1-560) directly, with the calculated Kd of 4.6 and 8.4 µM, respectively (Fig. S2B). Interestingly, in differential scanning fluorimetry (DSF) with recombinant CIP2A(1-560), only GNA exhibited a decrease in the melting temperature (Tm) indicative of the structural impact on CIP2A, and establishing functional relevance for high affinity GNA-CIP2A binding affinity (Fig. S2C).

**Figure 3:**
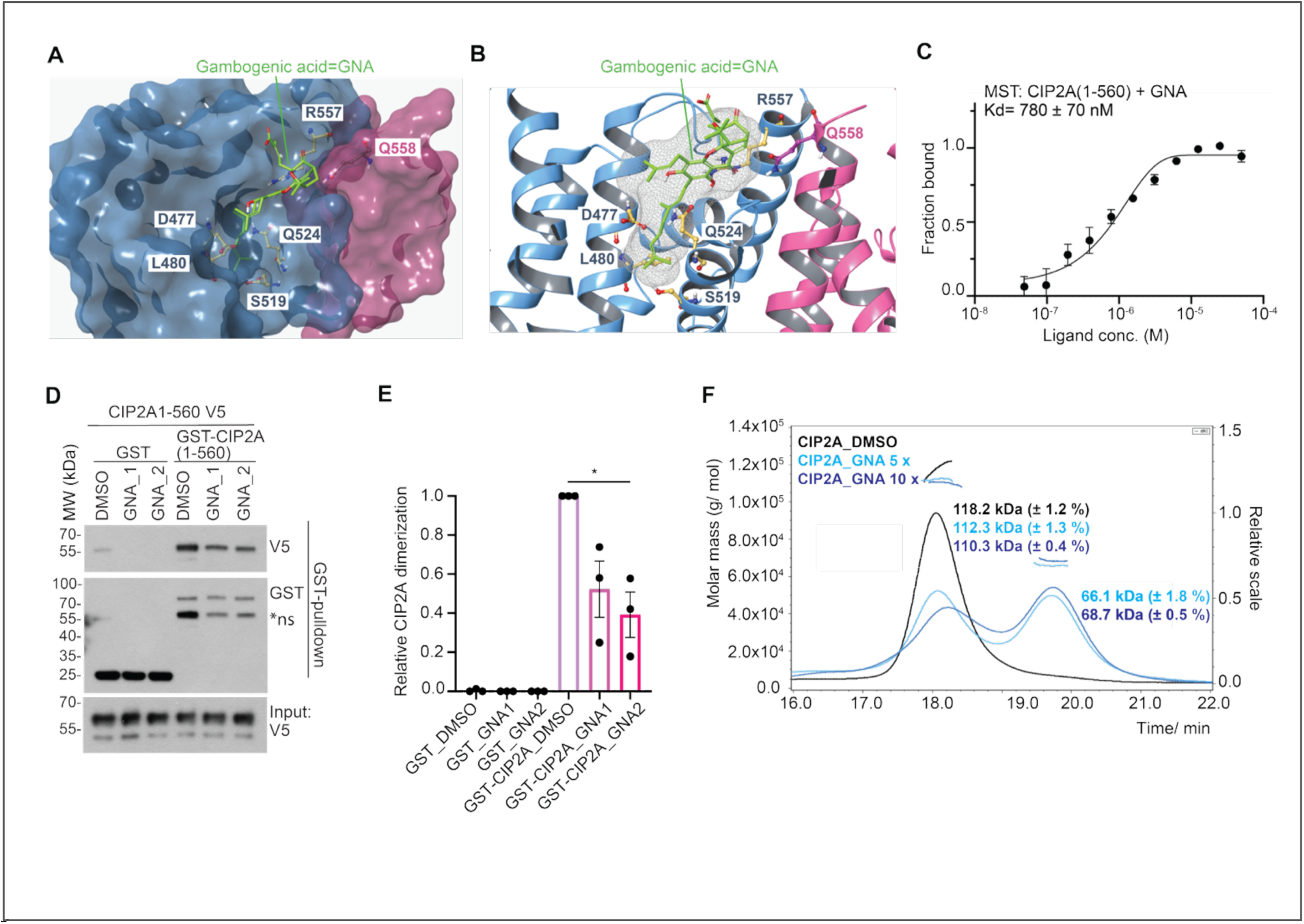
Direct binding and dimerization inhibitory activity on CIP2A by gambogenic acid. (**A,B**)) Gambogenic acid (GNA; sticks with green carbon atoms) occupies the binding pocket and extends toward the dimer interface, displacing Q558 (ball-and-stick with pink carbon atoms) and thereby perturbing dimerization (A) and ribbon (B) representations of Gambogenic acid (GNA) docked to the structure of CIP2A(1-560)(PDB: 5UFL), with individual CIP2A monomers colored in magenta and blue and GNA structure shown as green sticks with coloring based on element. The amino acids with modeled contacts with GNA are indicated including Q558 from the other monomer. (**C**) Microscale thermophoresis (MST) showing direct binding of gambogenic acid to CIP2A(1-560), with Kd = 780 nM ± 70 nM obtained from N = 3 biological repeats. (**D,E**) Representative blot of the GST-pull-down showing reduced CIP2A dimerization in presence of two different batches of gambogenic acid (GNA) used at 100 µM versus DMSO vesicle control (D), with quantified mean ± SEM of N = 3 biological repeats (E). (**F**) Size exclusion chromatography-coupled mass spectrometry multi-angle light scattering (SEC-MALS) analysis shows gambogenic acid (GNA) dissociates CIP2A(1-560) dimers. CIP2A(1-560) concentration was 7 µM and gambogenic acid concentration was 35 µM and 70 µM. Data collection and molar mass analyses were performed with Astra 7.1.4.

To experimentally corroborate GNA docking results, we performed the CIP2A dimerization assay using purified recombinant GST or V5-tagged CIP2A(1-560) fragments. To rule out the possible compound batch effect, we used two different batches of GNA, obtained from two independent manufacturers. Both batches of GNA were efficient in reducing CIP2A(1-560) dimerization *in vitro* by at least 50 % (Fig. 3D,E). Efficient inhibition of CIP2A(1-560) homodimerization was also observed with GIA, but not with DAP-19 (Fig. S3A,B). Further, yet another GNA derivative MAD28 (Yim et al., 2016) which did not interact with CIP2A by MST analysis (data not shown), also did not impact CIP2A homodimerization (Fig. S3C,D). The inhibitory effect of GNA on CIP2A homodimerization was confirmed by size-exclusion chromatography coupled multi-angle light scattering (SEC-MALS). Incubation of CIP2A(1-560) with five- or ten-fold molar excess of GNA triggered very efficient dimer dissociation in contrast to DMSO vehicle control (Fig. 3F).

To understand the structural basis why GNA and GIA, but not DAP-19, can break the CIP2A homodimerization, we compared their docking to the CIP2A pocket (Fig. S3E). GNA has a longer and more flexible hydrophobic tail than GIA or DAP-19, allowing it to better fit into the hydrophobic tunnel formed near the D477–Q524 interaction, and extend toward L480 (Fig. S3E). In contrast, the closed-pyrano ring structures of GIA and DAP-19 limit their conformational adaptability. DAP-19 also uniquely features a morpholine amide instead of the 2-methylbutenoic acid group, enabling an interaction with R557 via its carbonyl oxygen (Fig. S3E), which slightly shifts its position and prevents it from displacing Q558 of the opposing monomer. As a result, DAP-19 cannot disrupt CIP2A dimerization.

These data establish GNA and its derivatives as direct CIP2A binder molecules, and indicate that they disrupt CIP2A homodimerization by binding to the CIP2A pocket.

### Pocket region is required for impact of GNA on CIP2A homodimerization

To demonstrate that GNA engages with the CIP2A pocket region, we generated a biotinylated GNA variant (bGNA)(Fig. 4A). bGNA bound to recombinant CIP2A(1-560) based on MST analysis (Kd of 4.2 µM)(Fig. S4A). bGNA was next incubated either with GST-CIP2A(1-560), or with the N-terminal fragment GST-CIP2A(1-330), which does not contain pocket residues (Fig. 4A), and GNA binding to these CIP2A fragments was compared from streptavidin pull-down lysate by Western blotting. Indicative of crucial importance of the C-terminal region containing the CIP2A pocket for GNA-CIP2A interaction, GST-CIP2A(1-330) without the pocket region co-precipitated only at less than 20 % level of the GST-CIP2A(1-560) protein (Fig. 4B and 4C).

**Figure 4:**
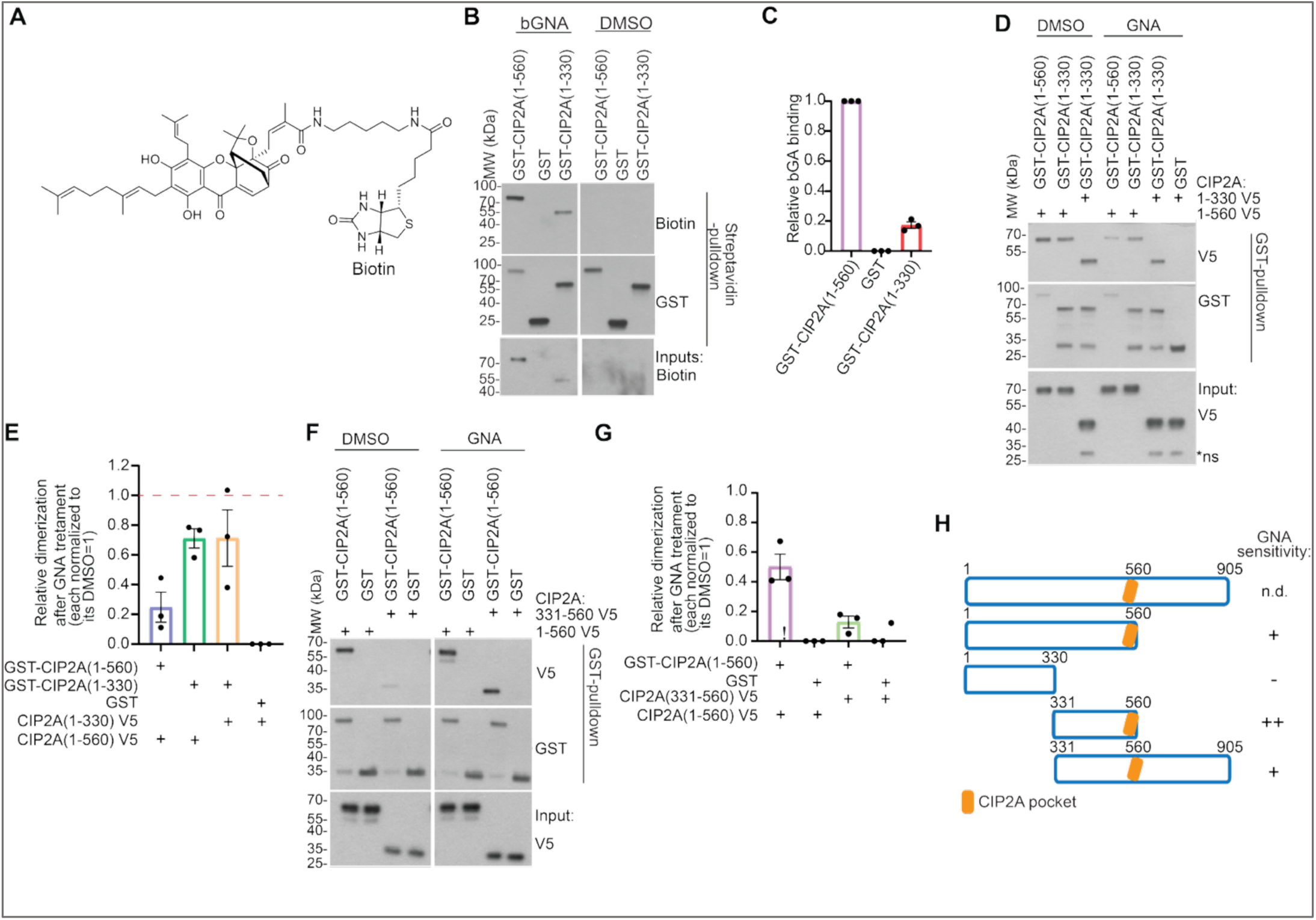
The pocket region is required for Gambogenic acid-elicited destabilization of CIP2A dimerization. (**A**) Chemical structure of biotinylated gambogenic acid (bGNA). (**B,C**) Representative blot of the streptavidin pull-down showing CIP2A(1-330) without pocket structure engages significantly less with biotinylated gambogenic acid versus CIP2A(1-560) (B)with quantified mean ± SEM of N = 3 biological repeats (C). (**D,E**) Representative blot of the GST pull-down analysis of impact of gambogenic acid (GNA) on CIP2A dimerization *in vitro* when comparing recombinant proteins containing the pocket region CIP2A(1-560) or not containing the pocket region (1-330) (D), with quantified mean ± SEM of N = 3 biological repeats (E). (**F,G**) Representative blot of the GST pull-down analysis of impact of N-terminal dimerization of CIP2A(1-560) on capacity of gambogenic acid (GNA) to inhibit dimerization of CIP2A (F), with quantified mean ± SEM of N = 3 biological repeats (G). (**H**) Schematic presentation of the CIP2A fragments used in pull-down assays, with CIP2A pocket indicated in orange and full-length CIP2A(1-905) shown as a reference. The relative impact of gambogenic acid (GNA) on CIP2A homodimerization of the fragments is indicated in a scale of - to ++.

While these results establish the region between amino acids 331-560 of CIP2A as the GNA binding region, we wanted to confirm that the impact of GNA on CIP2A homodimerization was also dependent on the pocket region. Upon the establishment of assays addressing this important point, we made an unprecedented observation that the N-terminal CIP2A(1-330) fragments, not containing the C-terminal dimerization interface, also co-precipitated in the GST pulldown assay (Fig. 4D). However, unlike CIP2A(1-560) homodimers, dimerization between CIP2A(1-330) fragments was not sensitive to disruption by GNA (Fig. 4D,E). Importantly, also the dimer between CIP2A(1-560) and CIP2A(1-330) was insensitive to GNA (Fig. 4D,E). However, the homodimer between CIP2A(1-560) and CIP2A(331-560), containing the entire pocket reaching from one monomer to another (Fig. 1B,C), but lacking the N-terminal interaction region from another dimerization partner, was found hypersensitive to GNA-elicited dimerization inhibition (Fig. 4F,G).

Finally, to address if GNA would be effective in disrupting CIP2A dimerization also in the context of the C-terminal tail of the protein, we produced a CIP2A fragment containing residues 331-905, spanning across pocket region, dimer interface, and the C-terminal tail of CIP2A. Upon dimerization assay with CIP2A(1-560)-V5, GNA could effectively disrupt CIP2A dimerization also with the GST-CIP2A(331-905) protein (Fig. S4B,C) indicating that the C-terminal tail does not interfere with binding of, or inhibitory effects on CIP2A by GNA.

As summarized in Figure 4H, these data establish requirement of the C-terminal pocket region mediating CIP2A homodimerization for GNA binding and its inhibitory impact on CIP2A homodimerization.

### Mutagenesis of the CIP2A pocket orifice impairs GNA-elicited CIP2A homodimerization inhibition

To further validate the pocket as the target of GNA activity in CIP2A homodimerization inhibition, we mutated two pocket amino acids L480 and S527 that based on modeling make direct contacts with GNA. Consequently, both amino acids were mutated to much bulkier tryptophans (L480W and S527W), as based on the structural analysis, the bulky substituent of these amino acids could hinder the optimal positioning of GNA (Fig. 5A). To test this theory, we generated these variants as GST-fusions and tested them in a dimerization assay against CIP2A(1-560) V5 WT variant. Whereas S527W mutation did not impact CIP2A(1-560) homodimerization, L480W mutation mimicked the GNA activity by efficiently inhibiting dimerization activity (Fig. 5B and 5C). Importantly, regardless of their spontaneous effects on homodimerization, both mutations effectively inhibited the ability of GNA to further break the homodimer (Fig. 5B and 5C). This can be more clearly appreciated from quantification, where the impact of each mutation is blotted against a DMSO-treated sample normalized to 1 (Fig. 5D). Whereas GNA caused approximately an 80 % reduction in WT CIP2A(1-560) dimerization, its impact against each of the pocket mutants was reduced to 40-50 % (Fig. 5D).

**Figure 5:**
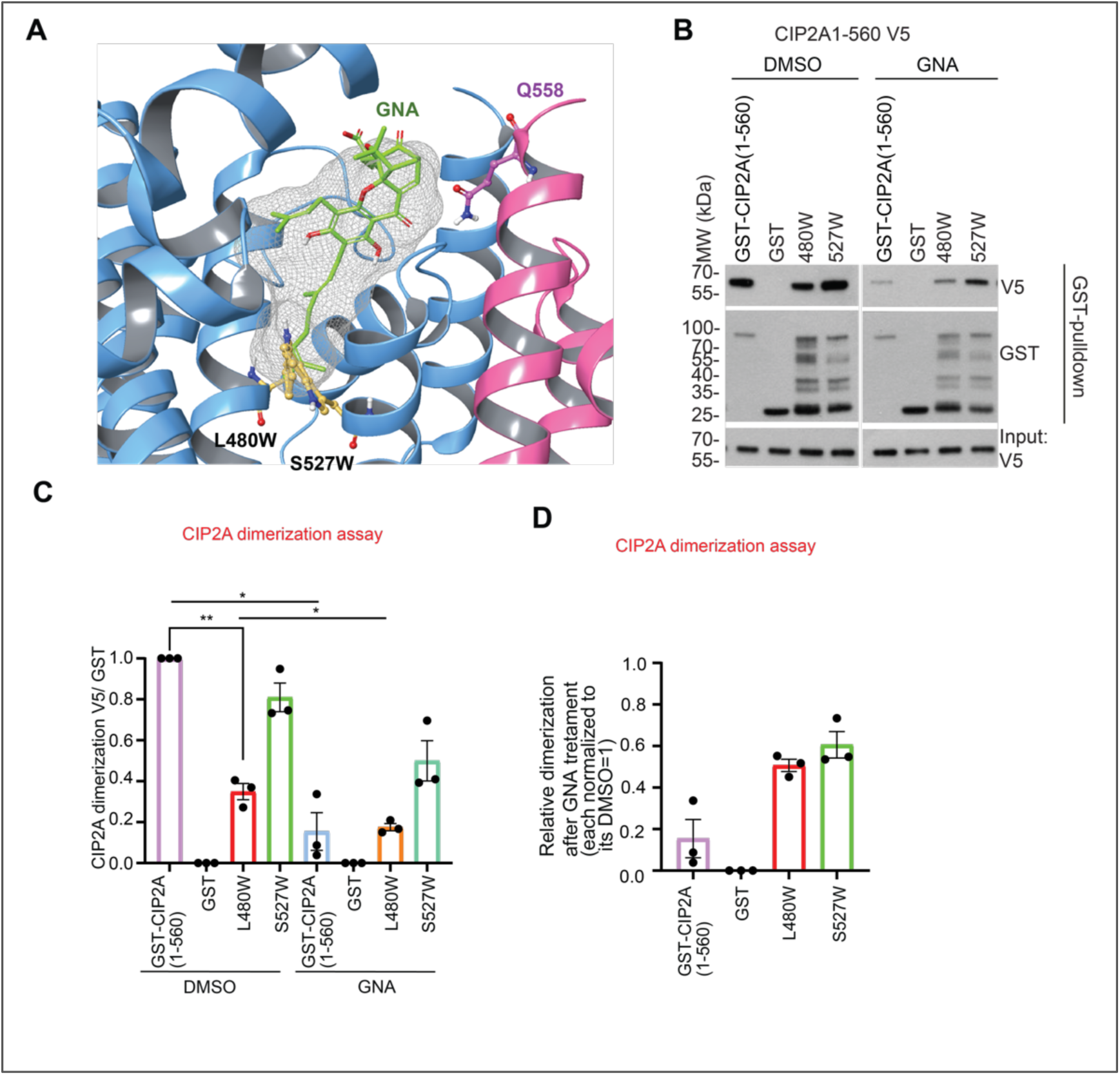
CIP2A pocket mutations inhibit gambogenic acid efficiency. (**A**) Substitution of L480 or S527 with tryptophan (orange ball-and-stick) is modelled to disrupt gambogenic acid (GNA) binding (sticks with green carbon atoms) by sterically occluding the spatial region occupied by the hydrophobic tail of GNA in the wild-type docking result. (**B,C**) Representative Western blot of the GST pull-down showing the impact of mutations on CIP2A homodimerization and on capacity of gambogenic acid (GNA) in disrupting CIP2A dimerization (B), with quantified mean ± SEM of N = 3 biological repeats all relative to WT CIP2A(1-560) (C). (**D**) Quantification demonstrating the relative impact of GNA on CIP2A homodimerization for either CIP2A WT, or each of the gate-keeper mutant CIP2A proteins normalized against dimerization efficiency of the same mutant treated with DMSO.

### CIP2A targeting by Gambogenic acid *in cellulo*

Although several alleged CIP2A inhibitors have been reported, the data demonstrating their target engagement in *cellular milieu* is lacking (Chen et al., 2023). To address if GNA would bind to CIP2A in *cellular milieu*, we incubated bGNA with cell lysates of MDA-MB-231 cells in which full-length V5-tagged CIP2A(1-905) was overexpressed and performed V5 pull-down. The cell lysates were used instead of living cells as GNA is known to trigger CIP2A degradation (Yu et al., 2016), and it may not have been possible to demonstrate its direct target engagement from living cells. Using biotin-HRP to detect bGNA that was physically associated with immunoprecipitated CIP2A(1-905)-V5, the experiments demonstrated repeatable and specific bGNA engagement with full-length CIP2A whereas no biotin signal was observed with any other conditions (Fig. 6A). Motivated by these results, we proceeded to validate the impact of GNA on endogenous CIP2A protein expression in cultured cancer cells. Treatment of triple-negative breast cancer cell lines MDA-MB-231 and MDA-MB-436, prostate cancer cell line 22RV1 and in HeLa cells with GNA inhibited CIP2A protein expression in 24 hours with concentrations that well align with measured GNA-CIP2A affinity *in vitro* (Fig. 6B and 3C). This same concentration range on GNA was sufficient also to inhibit cell viability of selected cell lines (Fig. S4D). Considering the importance of CIP2A for human breast cancer development and progression (Laine et al., 2021), we further examined the DepMap database (Arafeh, Shibue et al., 2025) for the supporting evidence. While no data for GNA was available, most breast cancer cell lines showed high sensitivity to GIA (Fig. 6C). However, due to previous studies demonstrating the multitargeting property of GNA and GIA in inhibiting oncogenicity (Zhou, Chen et al., 2025), we do not consider that their antioncogenic effects observed here can be attributed selectively to CIP2A inhibition.

**Figure 6:**
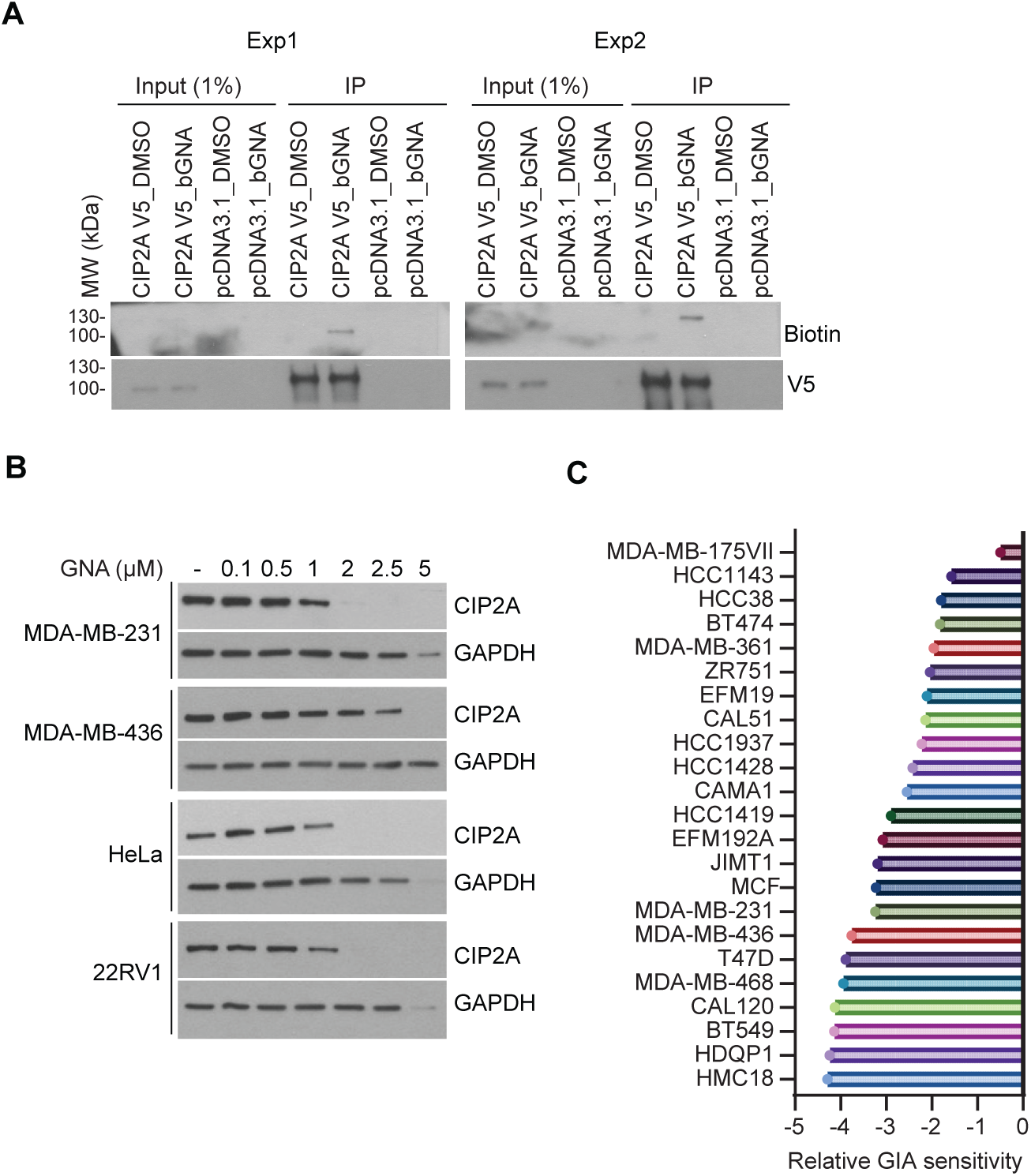
Gambogenic acid engages CIP2A *in cellulo*. (A) Western blot analysis of presence of biotinylated gambogenic acid (bGNA) V5-immunoprecipitation sample from the lysate of MDA-MB-231 cells transfected with in the CIP2A(1-905)V5 WT and incubated with bGNA. Blots from N = 2 independent biological repeats are shown. (B) Concentration dependent inhibition of endogenous CIP2A protein level by GNA in a panel of cancer cell lines. (C) Impact of Gambogic acid (GIA) on viability of breast cancer cell lines based on Dependency map data (PRISM Repurposing Public 24Q2 data at Depmap.org).

These data provide important validation of binding of GNA to full-length CIP2A in *cellular milieu* and its inhibitory effect on CIP2A protein expression. These finding are consistent with GNA binding to pocket, the protein destabilization effects of the CIP2A pocket mutations, and the impact of CIP2A homodimerization on its protein stability (Wang et al., 2017).

## Discussion

We discover here CIP2A pocket as a previously unknown structural vulnerability that determines CIP2A homodimerization and protein stability. This is a major advance in structural understanding of how central CIP2A functions are regulated, but also for assessment of small molecule druggability of CIP2A. Acknowledging the central role for CIP2A in pathogenesis of HRD cancers and in AD, we postulate that these results will help in development of targeted therapies for these disease types with clear unmet medical needs.

Based on both structural prediction programs, the CIP2A pocket extends over the dimerization interface (Fig. 1B,C). This indicates that CIP2A dimerization and pocket formation are functionally linked and explains why pocket mutations impact CIP2A homodimerization. The data published previously (Wang et al., 2017) and here (Fig. S1B), directly links CIP2A homodimerization propensity to protein stability *in cellulo,* but the flat CIP2A homodimerization interface does not exhibit any obvious druggability features. In fact, we tested a compound suggested to be able to bind to CIP2A homodimerization surface based on computational modeling (Bhowmick, Roy et al., 2022), but did not observe any inhibition of CIP2A homodimerization (Fig. S7 and S8). Thereby, identification of a pocket expanding to homodimerization interface, and impacting both homodimerization and the protein stability of CIP2A, constitutes a major step forward towards structure-based design of molecules interfering with pocket, and capable of reducing CIP2A protein expression to therapeutically beneficial levels.

In addition to providing significant novel structure-function understanding of CIP2A, we identify GNA and its derivatives as tool compounds that based on molecular modelling bind to CIP2A pocket. Use of these three highly related compounds (Fig. S2A) facilitated establishment of rudimentary structure-activity relationship correlating binding affinity with impact on protein folding, and CIP2A homodimerization. GNA and GIA also had anti-oncogenic activities in inhibition of CIP2A protein expression and cell viability across cell lines from different cancer types. GNA has been shown to target multiple oncogenic mechanisms beside CIP2A (Yim et al., 2016, Yu et al., 2016, Zhou et al., 2025). Thereby, our results are not to indicate that the effects of GNA and GIA on cancer cell viability could be attributed to CIP2A inhibition alone, and thus rescue experiments would not be meaningful. Instead, these results in combination are highly important as they demonstrate a structural feature on CIP2A via which its central functions including protein stability can be inhibited, and which can be targeted by small molecule compounds.

CIP2A is expressed at very low levels in most normal human tissues but gets overexpressed during the pathogenesis of most human cancer types and in AD (Chen et al., 2023, Hu et al., 2022, Shentu et al., 2018, Tang et al., 2018). Thereby the most reasonable strategy to target pathogenic functions of CIP2A is to reduce its protein expression, as is achieved here by targeting the CIP2A pocket (Fig. 2E-H). Protein de-stabilization is a particularly meaningful therapeutic strategy for CIP2A targeting in HRD cancers as it would target both oncogenic arms of CIP2A, PP2A inhibition arm driving tumorigenesis, and TopBP1 arm providing protection from mitotic cell death in BRCA mutant cells (Adam et al., 2021, Laine et al., 2021, Nagelli & Westermarck, 2024). Also the pathogenic effects of CIP2A-mediated PP2A inhibition in AD are related to its increased expression both in human AD samples and in mouse models of AD (Hu et al., 2022, Shentu et al., 2018). In these regards, our data identifying both a pocket on CIP2A amendable for small molecule binding, and CIP2A binder molecules, might facilitate development of targeted CIP2A degradation approaches for both disease classes based on pocket binders, or by proteolysis targeting chimera (PROTAC), and other related technologies (Luh, Scheib et al., 2020).

In summary, this study reveals novel structural insights into regulation of CIP2A homodimerization and protein stability, and proof-of-principle indicating small molecule druggability of the CIP2A pocket.

## Materials and Methods

### Cloning

CIP2A variants were generated by site-directed mutagenesis using polymerase chain reaction (PCR), with PfuUltra HF DNA polymerase (600380) or QuickChange Lightning Site-Directed Mutagenesis Kit (210518-5, both from Agilent Technologies) or Phusion Green Hot Start High-Fidelity PCR Master Mix (F-5665, Thermo Scientific) and DpnI enzyme from Quick Change Lightning Site-Directed Mutagenesis Kit (210518-5, Agilent Technologies).

For expression in *Escherichia coli*, the following CIP2A(1-560) variants were generated in pGEX4T1 vector, using GST-CIP2A(1-560) as a template and with the following primer pairs listed as forward and reverse and written in the 5’- 3’ direction: A476W (CTT GCT TGG GNAT GTA ATT TTG AAA ACT CTT GNAT TTG and CAA AAT TAC ACT CCA AGC AAG TTT GCA TAA TTC AGNA ATC), A516W (GCT TTT TGG TTA ACG TCA GNAT AAT AGNA GNAA CAA G and CGT TAA CCA AAA AGC CAA AGG AGT AAT CAA AC), L480W (GTA ATT TGG AAA ACT CTT GNAT TTG ATT AAC AAA C and G AGT TTT CCA AAT TAC ATC AGC AGC AAG TTT GC) and S527W (GTA CAG TGG GGNA CTG AGNA ATA TTA TTG GNAG G and CAGTCC CCA CTG TAC TTG TTC TCT ATT ATC TG).

For over-expression in mammalian cells, the following CIP2A variants were generated using the following primer pairs listed as forward and reverse and written in the 5’- 3’ direction: C472Y (GNAA TTA TAC AAA CTT GCT GNAT GTA ATT TTG and C AAG TTT GTA TAA TTC AGNA ATC TGC AAC CTT TG), K473E (GNAA TTA TGC GNAA CTT GCT GCT GNAT GTA ATT TTG and C AGC AAG TTC GCA TAA TTC AGNA ATC TGC AAC C), D484E (G AAA ACT CTT GNAA TTG ATT AAC AAA CTT AAA CCA TTG and GTT AAT CAA TTC AAG AGT TTT CAA AAT TAC ACT AGC), P494L (CA TTG GTT CTG GGT ATG GNAA GTA AGC TTC TAC and CAT ACC CAG AAC CAA TGG TTT AAG TTT GTT AAT C), R508H (GNAC CCA CAT TTG ATT ACT CCT TTG GCT TTT GC and GT AAT CAA ATG TGG GTC CTG AAG TAT TTT GTA G), T511A (GT TTG ATT GCG CCT TTG GCT TTT GCT TTA ACG and CAA AGG CGC AAT CAA ACG TGG GTC CTG AAG), T518M (GCT TTA ATG TCA GNAT AAT AGNA GNAA CAA GTA CAG and CT ATT ATC TGNA CAT TAA AGC AAA AGC CAA GGG AGT AAT C), S519K (GCT TTA ACG AAA GNAT AAT AGNA GNAA CAA GTA CAG TC and CT ATT ATC TTT CGT TAA AGC AAA AGC CAA AGG). A476W, L480W, A516W and S527W mutations were introduced using the same primer pairs as listed for the pGEX4T1 constructs. All plasmids for expression in mammalian cells were cloned in pcDNA3.1 vector using WT CIP2A(1-905) V5 His as a template. CIP2A(1-905) V5 S527W mutant was also generated using A476W mutant as a template. All constructs were verified by sequencing (Finnish Microarray and Sequencing Centre, Turku Bioscience Centre, Finland, and FIMM Helsinki, Finland).

For size-exclusion chromatography-coupled multi-angle light scattering (SEC-MALS) and differential scanning fluorimetry (DSF), CIP2A(1-560) was cloned into the pGTvL1-SGC vector (Structural Genomics Consortium, University of Oxford, UK) through ligation independent cloning method. For microscale thermophoresis (MST), CIP2A(1-560) with a C-terminal 6 x His-tag was synthesized by BioCat GmbH (Heidelberg, Germany) in a pUC57 cloning vector. CIP2A(1-560)-His was then cloned into a TEV site containing pGEX-4T-3 vector. The expression plasmids were verified by sequencing.

### Reagents

Gambogenic acid was from ALB Technology (#ALB-RS-5695) and from Chengdu Biopurity Phytochemicals Ltd (#BP2014). Gambogic acid, MAD-28 and DAP-19 were kindly provided by Dr. Leonard Neckers (National Cancer Institute, National Institutes of Health, Bethesda, MD) via Dr. Pia-Roos Mattjus (Åbo Akademi, Turku, Finland). AW00907 (CIP1) was from Maybridge screening library. Urea was from Sigma (#U5378), N-Lauroylsarcosine sodium salt (sarcozyl) was from Sigma (L9150), sodium phosphate monobasic (NaH_2_PO_4_) was from Sigma (#S3139), streptavidin agarose was from Thermo Fisher (#20347) and CellTiterGlo assay substrate (CTG) kit was from Promega (#G7571).

### Protein expression and purification for binding assays

For the *in vitro* binding assays, proteins were expressed and purified as described previously (Pavic et al., 2023). Represenative Coomassie stainings of the protein preps are shown in (Fig. S5). Standard protein expression scale was 3-4 L of LB medium. For expression in *Escherichia coli*, all CIP2A variants were cloned in pGEX vector, which produces proteins as thrombin-cleavable amino-terminal GST-fusion proteins. BL21 cells were used for over-expression. The overnight bacterial culture was inoculated in LB medium and incubated at 37°C until OD_600_ reached 0.6-0.9. Expression was induced with 0.2 mM isopropyl-β-D-1-thiogalactopyranoside (IPTG) for about 4 hat 23°C. The bacterial pellets were collected by centrifugation at 6,000 g at 4°C and stored at -20°C until purification. Cells were lysed by sonication on ice, in a buffer consisting of 200 mM Tris pH 8, 500 mM NaCl, 2 mM dithiothreitol (DTT), 0.5 % Tx-100, lysozyme (20 mg/ 150 mL) (Calbiochem 4403-1GM), and 1 x Pierce Protease Inhibitor Mini Tablets, EDTA-Free (Thermo Scientific A32955). Cleared lysate was incubated with glutathione agarose slurry (1:1 diluted with lysis buffer) at 4°C (from 4 h to overnight), with gentle rotation (Glutathione Sepharose 4B, 17-0756-01, GE Healthcare). Pelleted beads were washed extensively with washing buffer (same composition as lysis buffer, but without lysozyme), and then eluted using elution buffer consisting of 100 mM Tris pH 8, 200 mM NaCl, 5 mM DTT, 0.1 % Tx-100 and 20 mM Glutathione (L-Glutathione Reduced; Sigma-Aldrich G4251-5G). Samples were dialysed using Snakeskin MWCO 10k (Thermo Scientific 88243), or Slide-A-Lyzer Dialysis Cassette MWCO 10K (866383 Thermo Scientific), into a buffer containing 20 mM Tris pH 8, 150 mM NaCl, 2 mM DTT, 0.05 % Tx-100 and 10 % glycerol. If needed, the pooled fractions were concentrated using Amicon Ultra Centrifugal Filters (Merck Millipore), and concentration was determined by Coomassie staining (PageBlue Protein Staining Solution, Thermo Scientific 24630), using GST alone as internal standard.

B56α was also expressed as GST-fusion protein as described above. Affinity purification using glutathione agarose was conducted overnight at 4°C, with gentle agitation. Following extensive washes, GST tag was removed by overnight incubation at 4°C with AcTEV protease (Invitrogen 12575015). TEV was inactivated with phenylmethanesulfonyl fluoride (PMSF) at 1 mM final concentration, for 15 min at ambient temperature.

GST-CIP2A(331-905) was purified under denaturing conditions. First, the pellet was resuspended in a lysis buffer consisting of 100 mM Tris pH 8, 500 mM NaCl, 10 mM DTT, lysozyme (20 mg/ 150 mL lysis buffer, 1 x Pierce Protease Inhibitor Mini Tablets, EDTA-Free, 1 % SDS (w/ V) and 0.1 % sarkozyl (w/ V). The sample was subjected to sonication on ice until the solution turned clear and was then left to incubate on ice for 30 min to allow SDS precipitation. The lysate was cleared by centrifugation at 4°C and 13,000 rpm, for 20 min. Cleared lysate was incubated with 1 mL of glutathione agarose slurry overnight at 4°C with gentle rotation. Next day, the pelleted beads were rinsed extensively with washing buffer (the same composition as the lysis buffer, but without SDS and with 2 mM DTT) and eluted by adding 20 mM reduced glutathione to the washing buffer. The pellet left after the first sonication was resuspended in a buffer composed of 6 M urea, 10 mM DTT, 1 x Pierce Protease Inhibitor Mini Tablets, EDTA-Free, 0.1 M NaH_2_PO_4_ and 10 mM Tris pH 8 and subjected to the second round of sonication on ice. Cleared lysate was incubated with 1 mL of glutathione agarose slurry overnight at 4°C with gentle rotation.

### *In vitro* protein interaction assays

*In vitro* assays were performed as described previously (Pavic et al., 2023, Wang et al., 2017). All proteins were used at 10 pmol. Samples were diluted in the total volume of 150 µL of a reaction buffer consisting of 50 mM Tris HCl pH 7.5, 150 mM NaCl, 2 mM DTT, 0.2 % Igepal, 10 % glycerol, and incubated for 1 h at 37°C. Next, 5 µL input sample was withdrawn prior to adding 5 µL glutathione agarose (Glutathione Sepharose 4B,17-0756-01, GE Healthcare) (diluted 4 x in reaction buffer) to each sample. The samples were incubated for 1 h at ambient temperature, with moderate rotation. The beads were washed by adding 250 µL of reaction buffer for total of four buffer exchanges and for total of 1 h at 4°C, with moderate rotation. The bound complexes were eluted off the beads by adding 30 µL 2 x SDS-PAGE sample buffer and incubating for 10 min at 95°C. The eluted proteins were collected by centrifuging at 3,000 g for 1 min. Eluted materials were resolved on 4-20 % SDS-PAGE (Mini-Protean TGX Gels, Bio-Rad), transferred on PVDF membrane (Immobilon-P Transfer Membrane IPVH0010 from Merck Millipore) and blotted as indicated. Images were quantified using Image J. For each sample, the signal was divided with the signal obtained for the eluted GST bait protein. Next, the value for CIP2A(1-560) was set as one and the values for the CIP2A variants were calculated accordingly. Data was plotted using GraphPad Prism6.1, 10.2.0 and 10.4.2, showing mean + S.E.M. For reactions with the test compounds, a compound (or DMSO vehicle control) was pre-incubated with GST-tagged CIP2A variant for 30 min at 37°C, after which the V5-tagged CIP2A variant, or B56α, was added and the reaction proceeded for another hour at 37°C. The rest of the steps was as described above. All compounds were prepared in DMSO.

### CIP2A(1-560) recombinant protein expression and purification for characterization of binding to model compounds

For SEC-MALS and DSF, pGTvL1-SGC/ CIP2A(1-560) plasmid was transformed into *Escherichia coli* BL21 Gold cells. The protein was expressed as GST-fusion, in Terrific Broth (2.4 % w/v yeast extract, 1.2 % w/v tryptone, 0.5 % w/v glycerol, 0.017 M KH_2_PO_4_, 0.072 M K_2_HPO_4_) supplemented with 100 µg/ mL ampicillin. Expression was induced by the addition of IPTG to 0.4 mM final concentration at 18°C for 20 h. The cells were pelleted by centrifugation at 4,500 rcf for 20 min at 4°C. The pellets were re-suspended in lysis buffer (100 mM Tris, 100 mM NaCl, 20 % glycerol, 0.5 % CHAPS, 2 mM DTT, pH 8) and lysed by sonication (Sonoplus HD4100 Series, Bandelin; Sonoplus TS 113, Bandelin). Lysates were cleared by centrifugation at 35,000 x g for 30 min at 4°C. The GST fusion proteins were captured using Protino Glutathione Agarose 4B (Macherey-Nagel, Düren, Germany), and the GST was cleaved in situ at 4°C for 16 h using Tobacco Etch Virus (TEV) protease (Invitrogen, Life Technologies, Carlsbad, CA, USA). CIP2A was eluted using the following buffer PBS (10 mM Na_2_HPO4, 2 mM KH_2_PO4, 140 mM NaCl, 2.7 mM KCl, pH 7.4), 5 % glycerol, 2 mM DTT, pH 8. The TEV cleavage extended the CIP2A(1-560) construct by two additional N-terminal amino acid residues, M and S. The eluate containing CIP2A(1-560) was loaded on a HiLoad 16/60 Superdex 200 pg column (GE Healtcare) equilibrated with size exclusion (SEC) buffer (PBS, 0.03 % CHAPS, 2 mM DTT, pH 7.5). Fractions were analyzed by SDS-PAGE and those containing CIP2A(1-560) were pooled and concentrated to 1.5 mg/ mL using Amicon Ultra centrifugal devices (Merck Millipore, Burlington, MA, USA). For MST, CIP2A(1-560)-His was expressed and purified as described above except for lysate clarification (centrifugation at 15,000 x g for 30 min at 4°C), lysis buffer (100 mM Tris, 100 mM NaCl, 0.5 % CHAPS, 2 mM DTT, pH 8) and SEC buffer (1 x PBS, 0.03 % CHAPS, pH 7.5).

Size-exclusion chromatography-coupled multi-angle light scattering (SEC-MALS)SEC-MALS experiments were performed in-line with Äkta Pure system (GE Healtcare) and a MiniDAWN TREOS II dynamic light scattering detector and an Optilab T-rEX refractometer (Wyatt Technology). Samples were loaded on a Superdex 200 Increase 10/200GL (GE Healtcare) column and 1 X PBS, pH 7.5 was used as running buffer with a flow rate of 0.75 mL/ min. All experiments were performed at room temperature (RT). 90 μg of CIP2A(1-560) was mixed with 5 x or 10 x molar amount of GNA. Into CIP2A(1-560) alone sample, equivalent amount of DMSO (4 % v/v) was added. Samples were incubated at room temperature for 0, 15 or 30 min prior to injection. Data collection and analyses were performed with Astra 7.1.4 (Wyatt technologies). Protein concentration was determined from RI and UV using a dn/dc value of 0.185 mL/ g and an extinction coefficient of 0.368 mL/(mg.cm), respectively. Absolute molar mass was determined from LS using the Debye plot and Zimm’s model.

### Microscale thermophoresis

CIP2A(1-560) with a C-terminal 6 x His-tag was labelled with Monolith NT^TM^ Protein labelling kit RED-tris-NTA 2^nd^ Generation (Cat# MO-L018) according to the manufacturer’s instructions in assay buffer (50 mM Hepes, 300 mM NaCl, 0.03 % CHAPS, 0.05% Tween, 0.1 % PEG8000, pH 7.5). The measurements were performed using Monolith N.T. Automated Standard treated capillaries (MO-AK002). In the binding assays, CIP2A(1-560) concentration was kept constant at 50 nM, and 1:1 dilution series of compounds GNA, bGNA, GIA, and DAP19 were prepared in an assay buffer. The final concentration range of GNA and DAP19 was 100μM-49 nM. All samples contained 5 % (v/v) of DMSO. Measurements were performed in triplicates. The analysis was done using MO.Affinity Analysis v.2.2.7 and GraphPad Prism 10.4.1.

The data analyses were done using GraphPad Prims version 10.1.2 using One site – Total analysis to determine binding affinity and Dissociation – One phase exponential decay analysis.

### Differential Scanning Fluorometry

In the differential scanning fluorometry (DSF) experiments the final concentration of CIP2A(1-560) was 2.5 μM, and the concentrations of GNA, GIA, DAP19 and Biotinylated GNA were 25 μM. All samples were diluted into 1 x PBS, pH 7.5 0.03 % CHAPS, 5% DMSO, 5 x Sypro Orange (Invitrogen) in a total volume of 25 μL. Samples were incubated at RT for 15 min prior to measuring. Melt curves were measured using a CFX96 Real-Time PCR Detection System. Temperature increments 0.5°C/ 30 s from 20°C to 95°C were applied. All measurements were performed in triplicates. The fluorescence of each sample’s no-protein control was subtracted from the sample’s raw fluorescence data at each temperature. Subtracted replicates were averaged and then normalized from 0 to 1 and plotted as a function of temperature within the range of 20 - 55°C. Melting temperatures (T_m_) were determined at normalized 0.5 value after using GraphPad Prism’s (version 10.1.2) LOWESS Smoothing function adding 10 points between each 0.5°C increment. ΔT_m_ was calculated as following: ΔT_m_ = (T_m_ CIP2A DMSO) - (T_m_ CIP2A molecule).

### Antibodies and Western blotting

Mini-PROTEAN TGX TM Precast Protein Gels 4-20 % (BioRad) were routinely used for SDS-PAGE electrophoresis. After electrophoresis, sample were transferred to PVDF membrane by wet electro-blotting (200 mA, 75 min) and probed for specific antibodies, as indicated. The following antibodies were used: anti-V5 (monoclonal mouse (mM) Ab (E10/V4RR), MA5-15253), anti-V5 (mM Ab, R960-25), anti-GST (polyclonal rabbit (pR Ab), CAB4169) all from ThermoFisher Scientific, anti-CIP2A (mM Ab (2G10-3B5), sc-80659),, anti-GST (mM Ab (B-14), sc-138), anti-PR65 (pR Ab (H-300), sc-15355), anti-B56α (mM Ab (23), sc-136045), anti-β-Actin (mM Ab (C4), sc-47778), anti-c-Myc (mM Ab (9E10), sc-40), all from Santa Cruz Biotechnology, anti-PP2Ac (pR Ab, 2038S) and anti-Biotin HRP (mR Ab (D5A7), 5571S) both from Cell Signaling. andanti-GNAPDH (mM Ab (6C5), Hytest, 5G4-6C5).

Secondary antibodies used were: polyclonal goat anti-mouse immunoglobulin-HRP (P0447) and polyclonal swine anti-rabbit (P0399), both from Dako and used at 1:5,000 dilution for 1 H at ambient temperature. Pierce ECL Western Blotting Substrate (Thermo Scientific, 32106) was used for visualization by incubating with the membranes for 1 min and 1:1 dilution. The following films were used: Fuji Medical X-Ray Film (47410 19284, Fuji Film) and UltraCruz Autoradiography Film (sc-201697, Santa Cruz Biotechnology). Page Ruler Prestained Protein Ladder (26616 Thermo Scientific) was used as a protein size reference. Antibodies were diluted in TBS supplemented with 0.1 % Tween 20 and 5 % milk and were incubated overnight at 4°C or 1-3 H at room temperature.

### Cell culture

Cells were from ATCC and were cultured in a humidified incubator maintained at 37°C and 5 % CO_2_. 22RV1 and HCC-1937 cells were cultured in RPM1-1640 media (R5886-500ML, Sigma Life Science) supplemented with 10 % (V/V) fetal bovine serum (FBS), 0.5 % (V/V) penicillin/ streptomycin (10,000 U/ 10 mg per mL, Sigma) and 2 mM L-Glutamine (Biowest). MDA-MB-468 cells were cultured in RPMI supplemented as above, but with 5 % FBS. MDA-MB-231 and MDA-MB-436 cells were maintained in DMEM (D6171-500ML, Sigma Life Science) and supplemented as above. Cells were routinely passaged 2-3 times per week and regularly tested for mycoplasma.

### Analysis of protein expression of CIP2A(1-905) V5 variants in mammalian cells

Cells were seeded in 12 well-plates and transfected using Lipofectamine 3000 or jetPRIME (Polyplus, 101000046) transfecting reagents at 3:1 and 2:1 DNA:transfecting reagent ratios (µL of transfecting reagent per µg of DNA transfected). The samples were collected 24 h post-transfection. 22RV1 cells were scraped on PBS, mixed with 2 x SDS-PAGE sample buffer and incubated at 95°C for 10 min. MDA-MB-231 cells were collected on lysis buffer consisting of 20 mM Tris pH 8, 150 mM NaCl, 2 mM DTT, 0.05 % Tx-100 and 1 x Pierce Protease Inhibitor Mini Tablets, EDTA-Free and lysed by sonication on ice for 2.5 min. Clarified lysates were loaded on 4-20 % SDS-PAGE gels and analysed by Western blotting using specific antibodies. Images were quantified using Image J. For quantification, for each sample, the signal from the V5 antibody was first normalized against β-Actin or GNAPDH. The value for WT CIP2A(1-905) V5 was set as one and the values for the CIP2A mutants were calculated accordingly. Graphs were plotted using GraphPad Prism6.1 and 10.2.0.

### Synthesis and characterization of biotinylated gambogenic acid (bGNA)

GNA (3.5 mg, 5.6 µmol) and EZ-link (5-biotinamido) pentylamine (2.0 mg, 6.1 µmol) were dissolved in DMSO (250 µL) followed by addition of benzotriazol-1-yloxy-tris(1-pyrrolidinyl)phosphonium hexafluorophosphate (PyBOP, 3.2 mg, 6.1 µmol) and diisopropylethylamine (DIEA, 2.0 µl, 11.1 µmol). The reaction was allowed to proceed overnight at RT. The reaction was stopped by adding ethyl acetate and water. The organic phase was collected and the aqueous phase was extracted two times with ethyl acetate. The crude material was purified by RP-HPLC with Phenomemex 250 × 10 Kinetex^TM^ 5 µm C18 100 Å column and a linear gradient of acetonitrile (30 – 80 %). The collected fractions were lyophilized affording the biotinGNA(4.5 mg, 86 % yield). The structure of the bGNA and the ^1^H and ^13^C NMR data of the product is shown in supplementary figures S6.

### Biotinylated gambogenic acid in pull-down assays

For pull-down assays using purified recombinant proteins, bGNA (50 µM) was incubated with CIP2A variants (10 pmol) for 30 min at 37°C, followed by precipitation of the formed complexes for 1 h at room temperature. The beads were rinsed two to three times for the total of 30 min at room temperature and with mild rotation. Bound material was eluted off the beads by adding 30 µL 2 x SDS-PAGE sample buffer and incubating 10 min at 95°C, resolved by SDS-PAGE and analyzed by Western blotting. For pull-downs from MDA-MB-231 cell lysate, 10 µg of cDNA encoding pcDNA3.1 empty vector control or CIP2A(1-905)V5 was transfected into 10 cm dish using JetPrime. Cells were collected about 48 h after transfection on 1 mL lysis buffer consisting of 20 mM Tris pH 8, 150 mM NaCl, 2 mM DTT, 0.05 % Tx-100 and 1 x Pierce Protease Inhibitor Mini Tablets, EDTA-Free. From the cleared lysate, 10 µL sample (1 % Input) was withheld for SDS-PAGE and to the remains bGNA was added at the final concentration of 100 µM. The samples were incubated for 1 h at 4°C, then 20 µL of anti-V5 agarose affinity gel produced in mice (A7345-1ML, Sigma) (40 µL slurry) was added to each sample and incubated for additional 3 h. The beads were rinsed twice with 500 µL of lysis buffer and eluted by applying 30 µL of 2 x SDS-PAGE sample buffer and incubating at 95°C. The samples were resolved on 4-20 % SDS-PAGE gels and analyzed by Western blotting using specific antibodies.

### Flexible molecular docking

The crystal structure of CIP2A (PDB ID: 5UFL) was retrieved from the Protein Database (Berman, Westbrook et al., 2000, ww, 2019). The structure was protonated using REDUCE v3.24 (Word, Lovell et al., 1999) to assign hydrogen atoms appropriately. All crystallographic water molecules were removed prior to further modeling procedures. The structures of Gambogenic Acid (GNA), Gambogic Acid (GIA), and DAP-19 were prepared using the LigPrep module in MAESTRO2025-1 (Schrödinger, LLC, New York, NY, 2025) with the OPLS3 force field (Harder, Damm et al., 2016). The ionization states were generated at physiological pH 7.4 to ensure biologically relevant protonation. To identify potential ligand-binding cavities, Panther (Niinivehmas et al., 2015) was utilized for structure-based cavity detection. The identified binding sites were further refined and optimized. Subsequently, ShaEP (Vainio et al., 2009) was used to superimpose the known ligand GNA onto the Panther-derived models to align and validate the binding site geometry. The resulting composite site was used for subsequent molecular docking studies. Flexible molecular docking was carried out using PLANTS v1.2 (Korb, Stutzle et al., 2009). A docking box with a 12 Å radius was defined, centered on the geometric midpoint of the key binding site residues R557, D477, K473, and E527.

### Molecular Dynamics (MD) simulation

The molecular dynamics (MD) simulations were performed using the AMBER24 package (Case, Aktulga et al., 2024). The ff14SB force field (Maier, Martinez et al., 2015) was applied for the protein parameters, while ligands were parametrized using the General AMBER force field (Wang, Wolf et al., 2004). The protein-ligand complex was solvated in a cubic TIP3P water box (Jorgensen, Chandrasekhar et al., 1983), extending 15 Å beyond the protein atoms in all directions. Counter ions (Cl⁻) were added to neutralize the overall system charge.

Three independent MD simulation runs were performed for each studied system. Energy minimization was carried out in four stages using the conjugate gradient method: (a) 2,000 steps with all atoms except hydrogen restrained (1 kcal/ mol); (b) 2,000 steps with all atoms except hydrogen and solvent molecules restrained; (c) 2,000 steps with restraints only on the protein backbone; and (d) 2,000 steps with no restraints. After energy minimization, the system was gradually heated to 310 K with a 4 kcal/ mol restraint on the protein backbone. A restrained equilibration simulation of 2 ns was then performed, during which the restraint force was reduced stepwise every 250 ps. Subsequently, a 50-ns production simulation was conducted without restraints, using a 2-fs time step under the NPT ensemble at 1 atm. Periodic boundary conditions were applied, with a 10 Å cutoff for non-bonded interactions. Long-range electrostatic interactions were treated using the particle mesh Ewald (PME) method (Darden, York et al., 1993, Essmann, Perera et al., 1995). The figures were generated using MAESTRO2025-1 (Schrödinger) and the MD simulation trajectories were examined using VMD1.9.4a12 (Humphrey, Dalke et al., 1996).

### Statistics and Data Analysis

Data were analyzed using GraphPad Prism10.4.2. The number of independent biological repeats is indicated in the figure legends. Plotted are mean with standard error (SEM) unless written otherwise. Western blot experiments were analyzed using unpaired t-test with Welch’s correction. A p-value of < 0.05 was considered statistically significant. SEC-MALS data analyses were done with GraphPad Prims version 10.1.2 using One site – Total analysis to determine binding affinity and Dissociation – One phase exponential decay analysis. *=p<0.05, **=p<0.01, ***=p<0.001, ****=p<0.0001.

## Acknowledgements

Gambogic acid, MAD-28 and DAP-19 were kindly provided by Dr. Leonard Neckers (National Cancer Institute, National Institutes of Health, Bethesda, MD) via Dr. Pia-Roos Mattjus (Åbo Akademi, Turku, Finland). We thank Henrik Honkanen for experimental data related to effect of GNA on cell viability and Mrs Taina Kalevo-Mattila for expert technical assistance. This research was funded by Sigrid Juselius Foundation (JW), Cancer Foundation Finland (JW), Magnus Enrooth Foundation (KP), Albin Johanssons Foundation (KP), Finnish Cultural Foundation (KP), Novo Nordisk Foundation (OTP, (0075825, 0075825, 0096035; and UP (0076296)), Research Council of Finland (UP 283481); Doctoral Programme on Drug Research and Diagnostics (PR); National Doctoral Education Pilot Based on the Immune System (PM-G). The Finnish IT Center for Science (CSC) is acknowledged for generous computational resources (OTP: Project Nos. 2002811, jyy2516 and jyy2585).

## Disclosure and competing interests statement

OTP is a founder and shareholder of Aurlide Ltd. Other authors declare no competing interests.

## Supplementary Figure Legends

**Figure S1:**
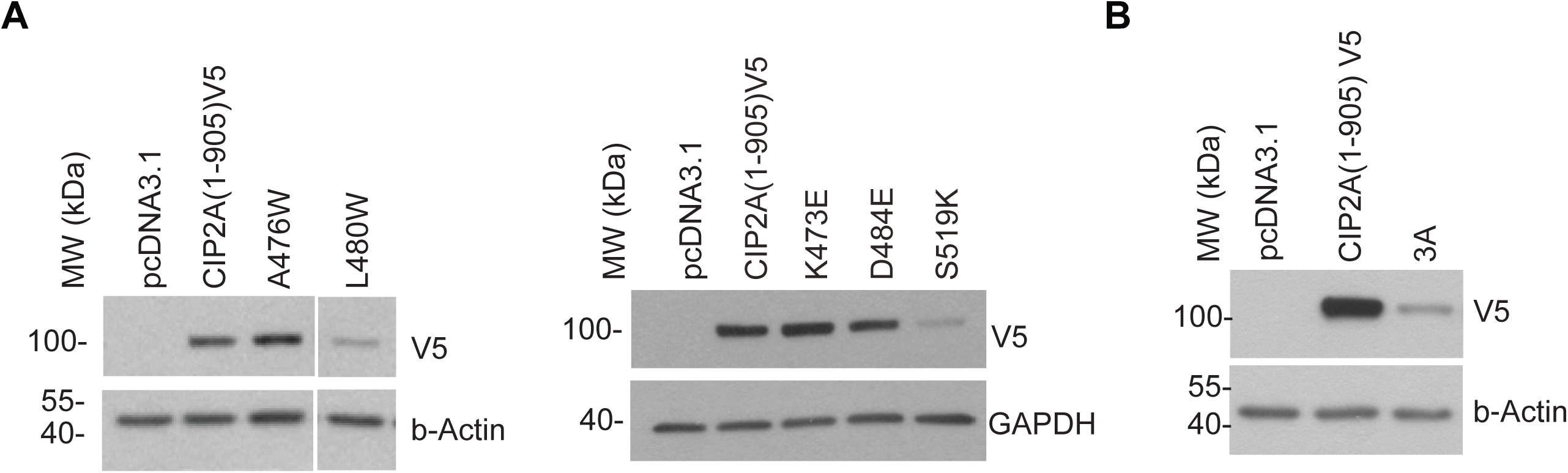
Representative blot showing protein expression level of CIP2A mutations predicted from the modeling (K473E, D484E, S519K) (C), with quantified mean ± SEM of N = 3 biological repeats (D).

**Figure S2:**
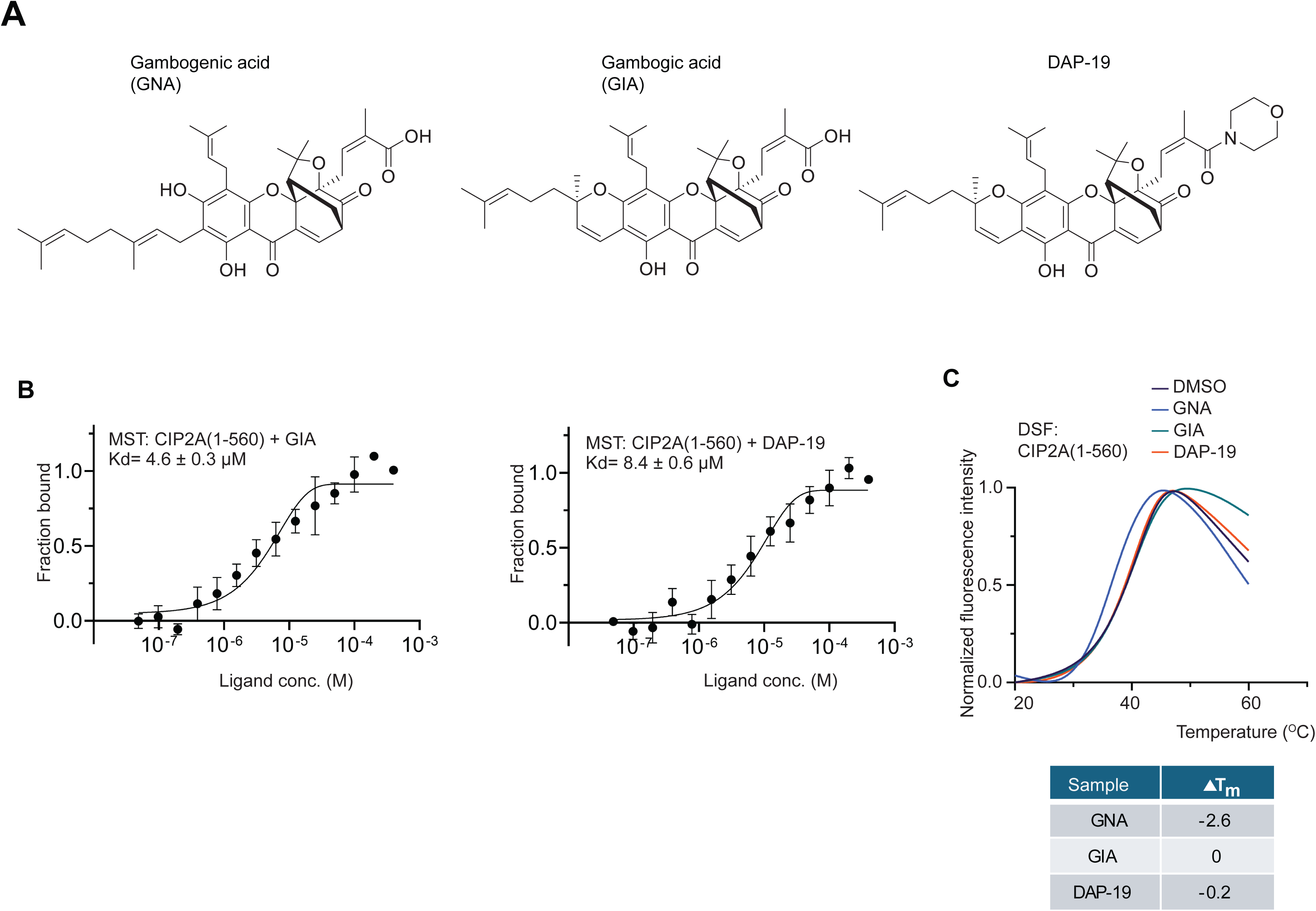
(**A**) Chemical structures of gambogenic acid (GNA), gambogic acid (GIA) and DAP-19. (**B**) Direct binding of gambogic acid (GIA) and DAP-19 to CIP2A(1-560) using microscale thermophoresis (MST). N = 3 biological repeats. (**C**) Impact of gambogenic acid (GNA), gambogic acid (GIA) and DAP-19 on CIP2A(1-560) thermostability using differential scanning fluorimetry (DSF). Indicated changes in the protein melting temperature from triplicate measurements were obtained using a CFX96 Real-Time PCR Detection System. Normalized melt curves were plotted and melting temperatures analyzed with GraphPad Prism 10.4.1.

**Figure S3:**
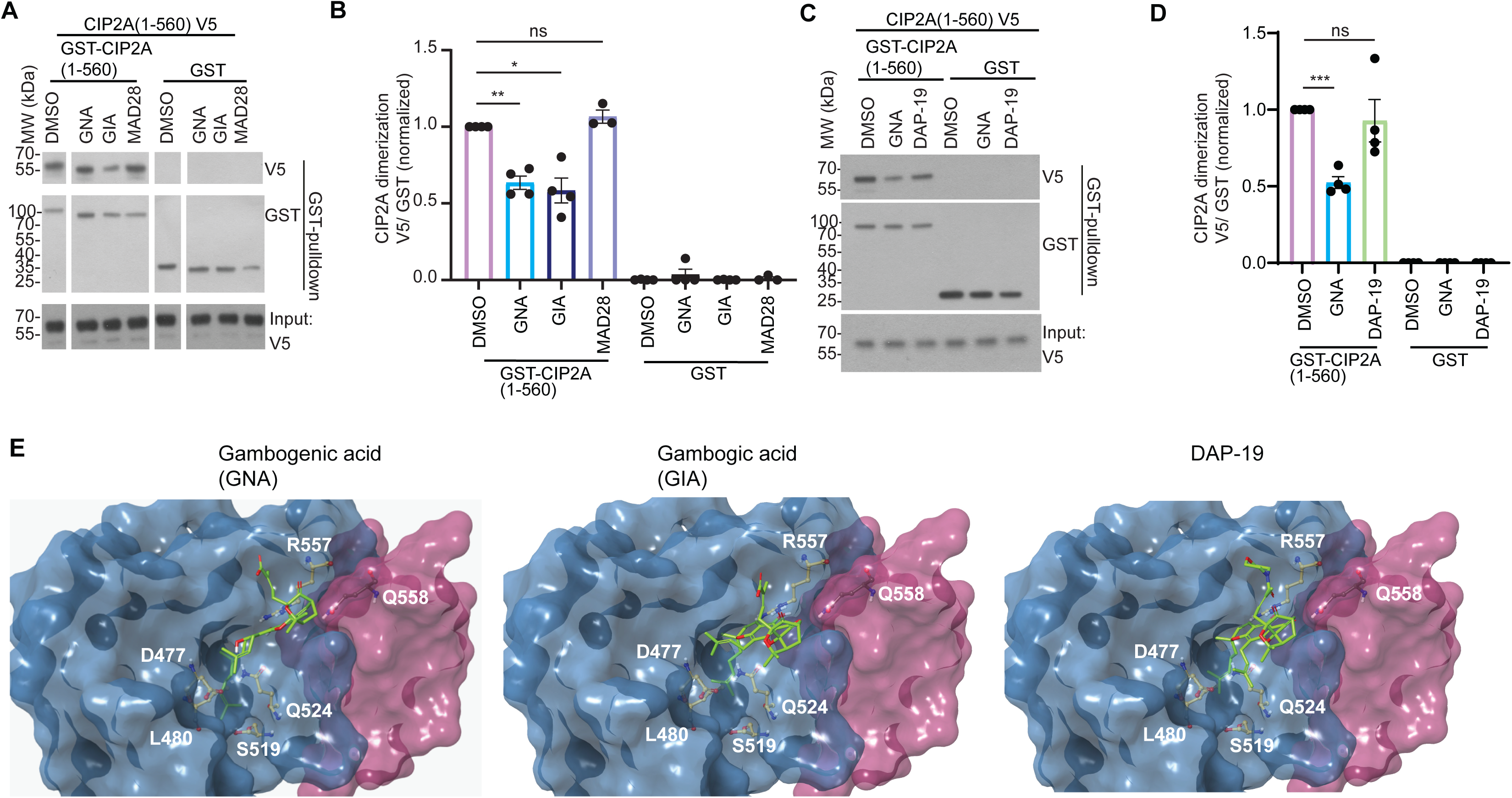
(**A,B**) Representative blot of the GST pull-down showing CIP2A dimerization disrupting effects of gambogic acid (GIA) and MAD-28 in contrast to gambogenic acid (GNA), all at 50 µM, using purified recombinant proteins *in vitro* (A), with quantified mean ± SEM of N = 4 biological repeats (B). (**C,D**) Representative blot of the GST pull-down showing CIP2A dimerization disrupting effects of DAP-19 in contrast to gambogenic acid (GNA), both at 50 µM, using purified recombinant proteins *in vitro* (A), with quantified mean ± SEM of N = 4 biological repeats (B). (**E**) Predicted binding modes of gambogenic acid (GNA), gambogic acid (GIA) and DAP-19 to CIP2A pocket based on molecular dynamics simulations.

**Figure S4:**
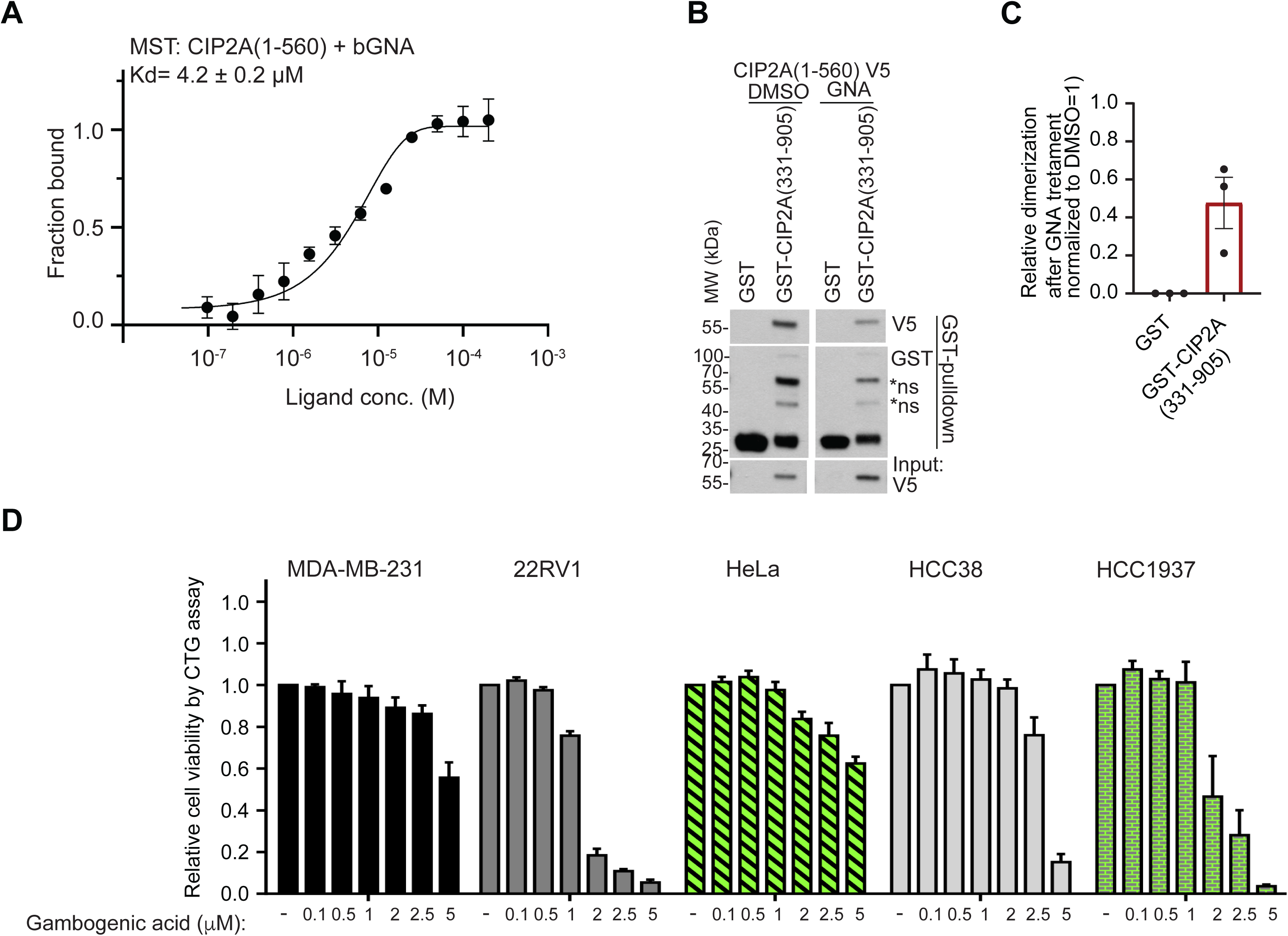
(**A**) Microscale thermophoresis (MST) showing direct binding of biotinylated gambogenic acid to CIP2A(1-560). N = 3 biological repeats. (**B,C**) Representative blot of the GST pull-down showing gambogenic acid reduces CIP2A dimerization *in vitro* in the presence of C-terminal CIP2A tail (residues 331-905) (B), with quantified mean ± SEM of N = 3 biological repeats (C). (**D**) Impact of gambogenic acid on cell viability of indicated cell lines as measured by Cell Titer Glow (CTG) assay 24 hours after treatment. Shown is mean + S.D from four technical replicate samples.

**Figure S5.**
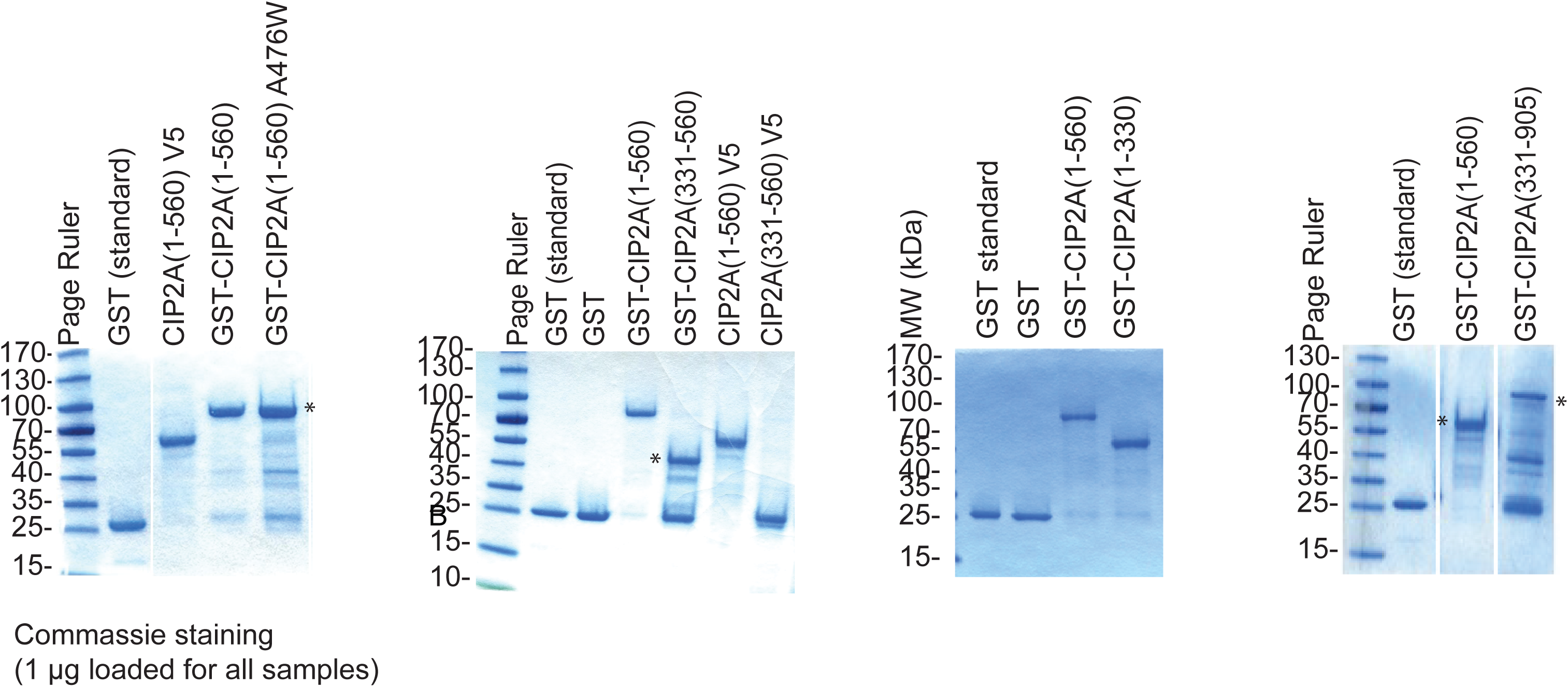
Coomassie staining of the purified recombinant CIP2A fragments expressed in *Escherichia coli*, used in *in vitro* protein interaction assays, with 1 µg loaded and GST alone used as internal concentration reference. Star indicates expected MW.

**Figure S6.**
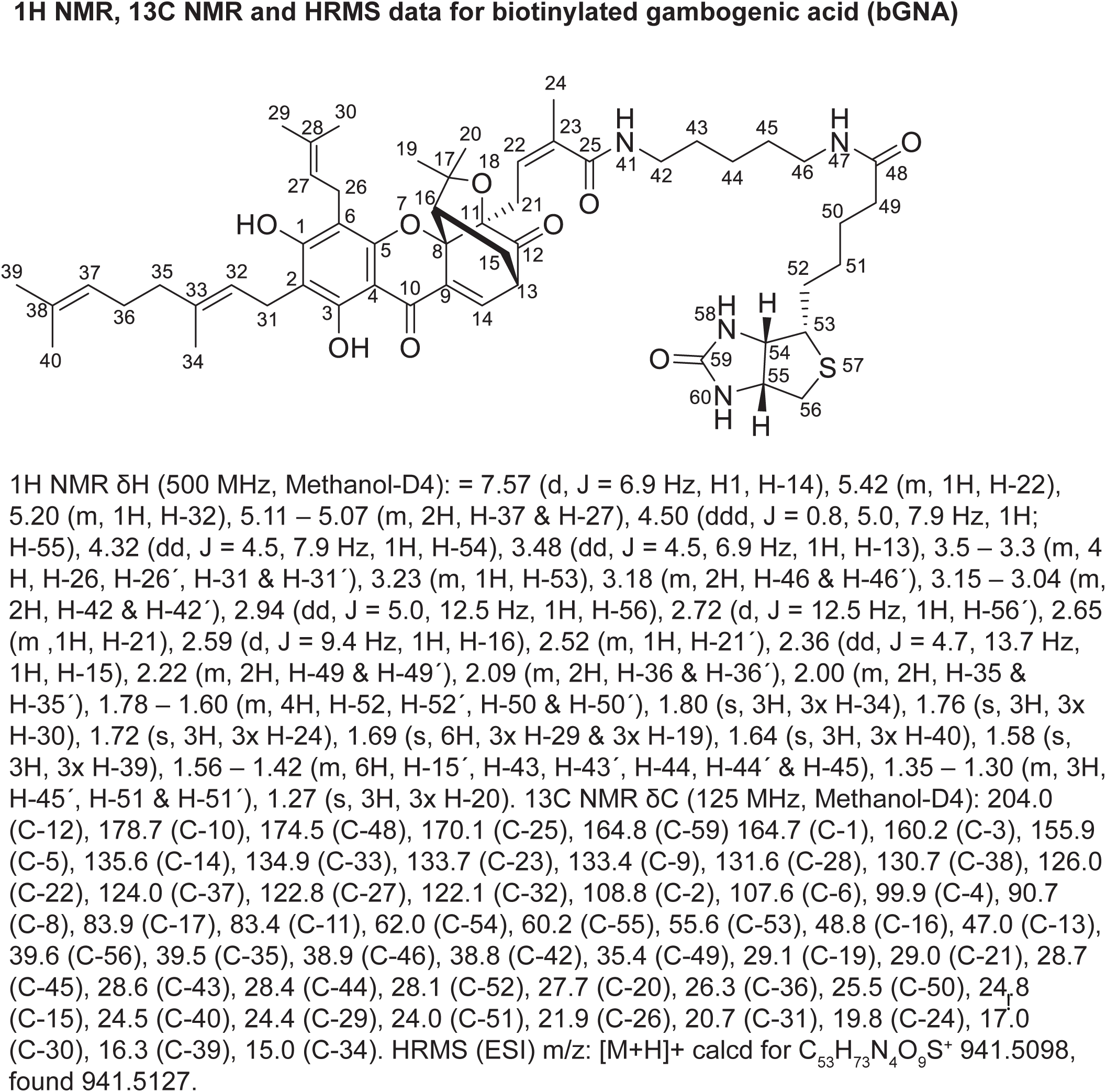
Structure and NMR coordinates of the biotinylated GNA (bGNA).

**Figure S7.**
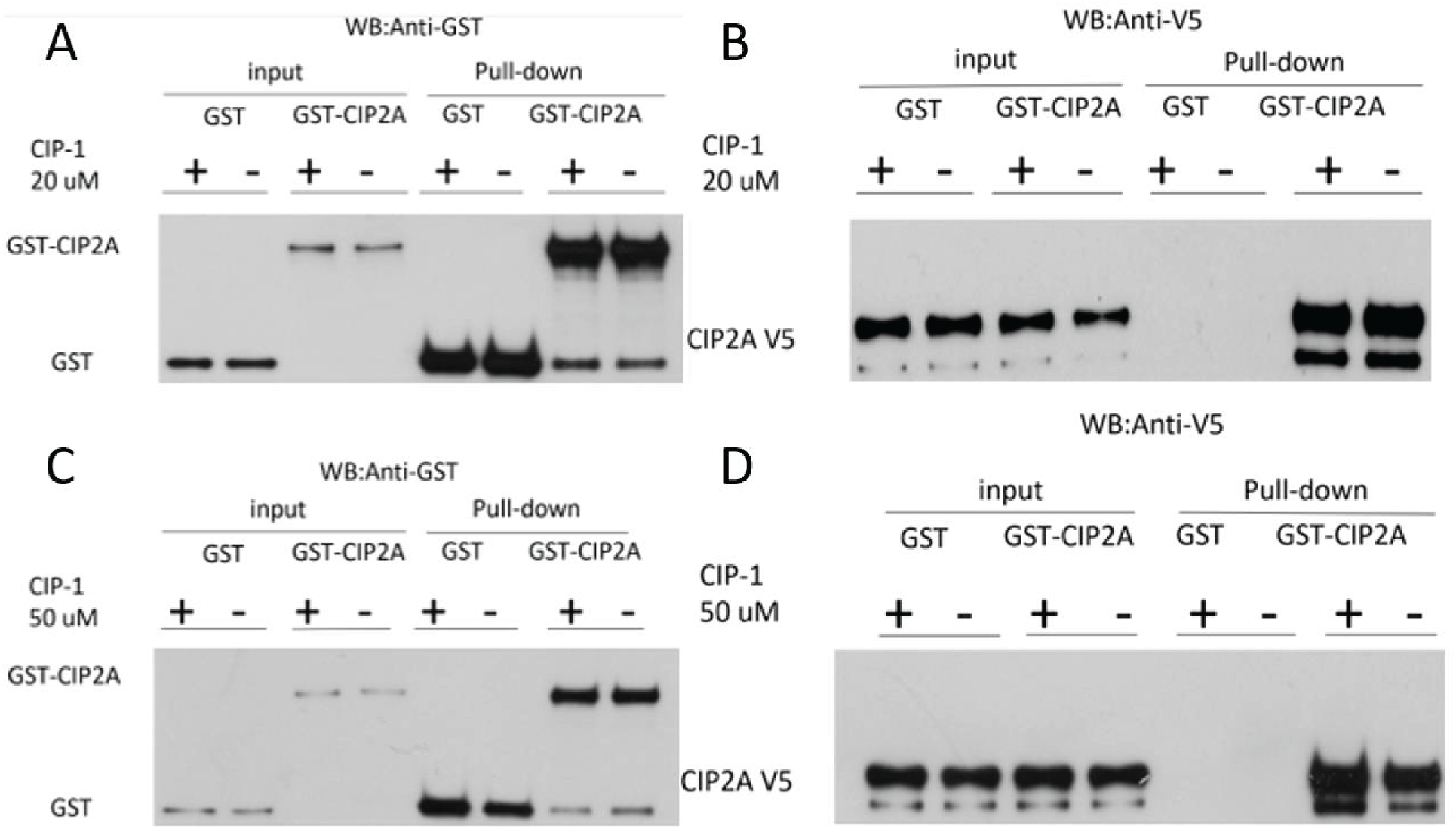
GST-pulldown assay testing the effect of AW00907 (CIP1) on CIP2A (1-560) homodimerization. Equal molar amounts of GST and GST-CIP2A(1-560)were incubated with CIP2A(1-560)-V5 fragment for 1h at 37 Celsius degree in the presence or absence of either 20 uM (A and C) or 50 uM (B and D) of CIP1.

**Figure S8.**
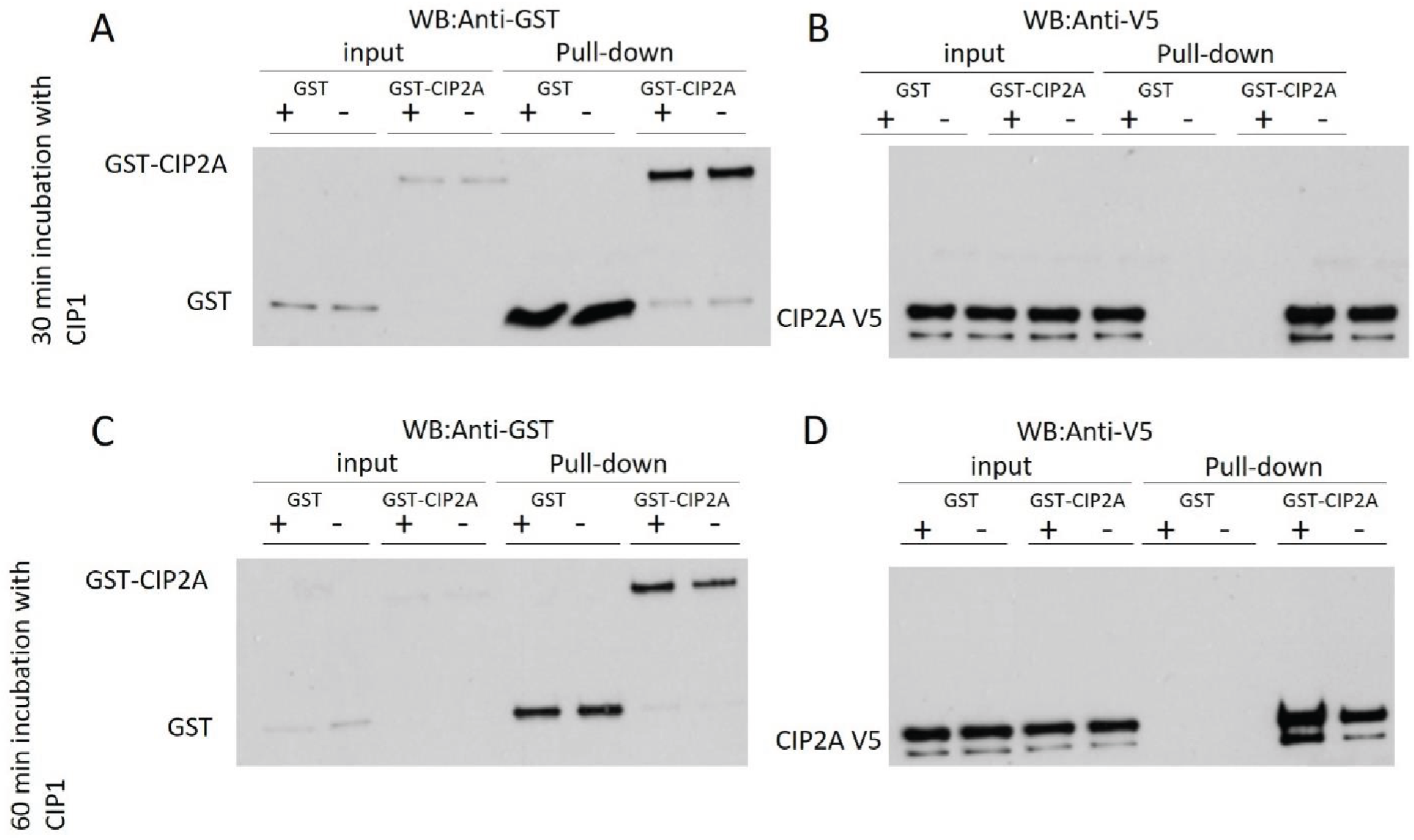
GST-pulldown assay testing the effect of 100 uM of AW00907 (CIP1) on CIP2A (1-560) homodimerization. Equal molar amounts of GST and GST-CIP2A(1-560)were incubated with CIP2A(1-560)-V5 fragment and with or without CIP1 for either 30 min (A and B) or 60 min (C and D).

